# Sequential Molecular Interactions Shape Aβ42 Aggregation, Propagation, and Toxicity

**DOI:** 10.64898/2026.08.27.747468

**Authors:** Jofre Seira Curto, Genís Pérez Collell, Marina Romero Ruiz, Sandra Villegas Hernandez, Maria Rosario Fernandez, Natalia Sanchez de Groot

## Abstract

Protein aggregation is a context-dependent process in which the molecular environment can influence the properties of the resulting assemblies. In biological systems, these interactions can occur sequentially, as aggregates formed in one cellular or tissue context may encounter different molecular partners and act as seeds in subsequent aggregation events. Here, we used sequential seeding as a controlled experimental model of this temporal and contextual complexity to investigate how prion-like sequences from the gut microbiome modulate amyloid-β aggregation across successive aggregation cycles. Combining kinetic, biophysical, conformational, and toxicity analyses, we show that early interactions with exogenous peptides modify the properties of first-generation Aβ40- and Aβ42-derived seeds, resulting in propagated Aβ42 assemblies with distinct molecular and functional properties. These findings support an Interaction History model in which exogenous sequences bias the emergence of aggregate populations whose properties and subsequent propagation depend on the molecular contexts experienced during earlier aggregation events. Overall, our results present Aβ aggregation as a history-dependent process and suggest that single-step assays may fail to capture aggregate diversity that emerges across successive aggregation cycles.

## 1. Introduction

Protein aggregation is a dynamic self-assembly process in which multiple conformational states can emerge depending on the molecular environment and assembly pathway. In Alzheimer’s disease (AD), extracellular aggregation of amyloid-beta (Aβ) peptides leads to the formation of the characteristic amyloid plaques [1–5]. Aβ exists mainly in two isoforms: Aβ40, the most abundant, and Aβ42, which is more aggregation-prone and predominantly found in amyloid plaques [4,6] (Figure 1). These isoforms can interact during fibrillization, modulating each other’s aggregation behavior [2,7–11]. Consistent with this, overexpression of Aβ40 has been reported to suppress Aβ42 deposition, suggesting a regulatory or protective role [9,12,13]. Additionally, analyses of patient-derived samples have shown that Aβ40 and Aβ42 can co-aggregate into interlaced fibrils [9,14,15].

**Figure 1.**
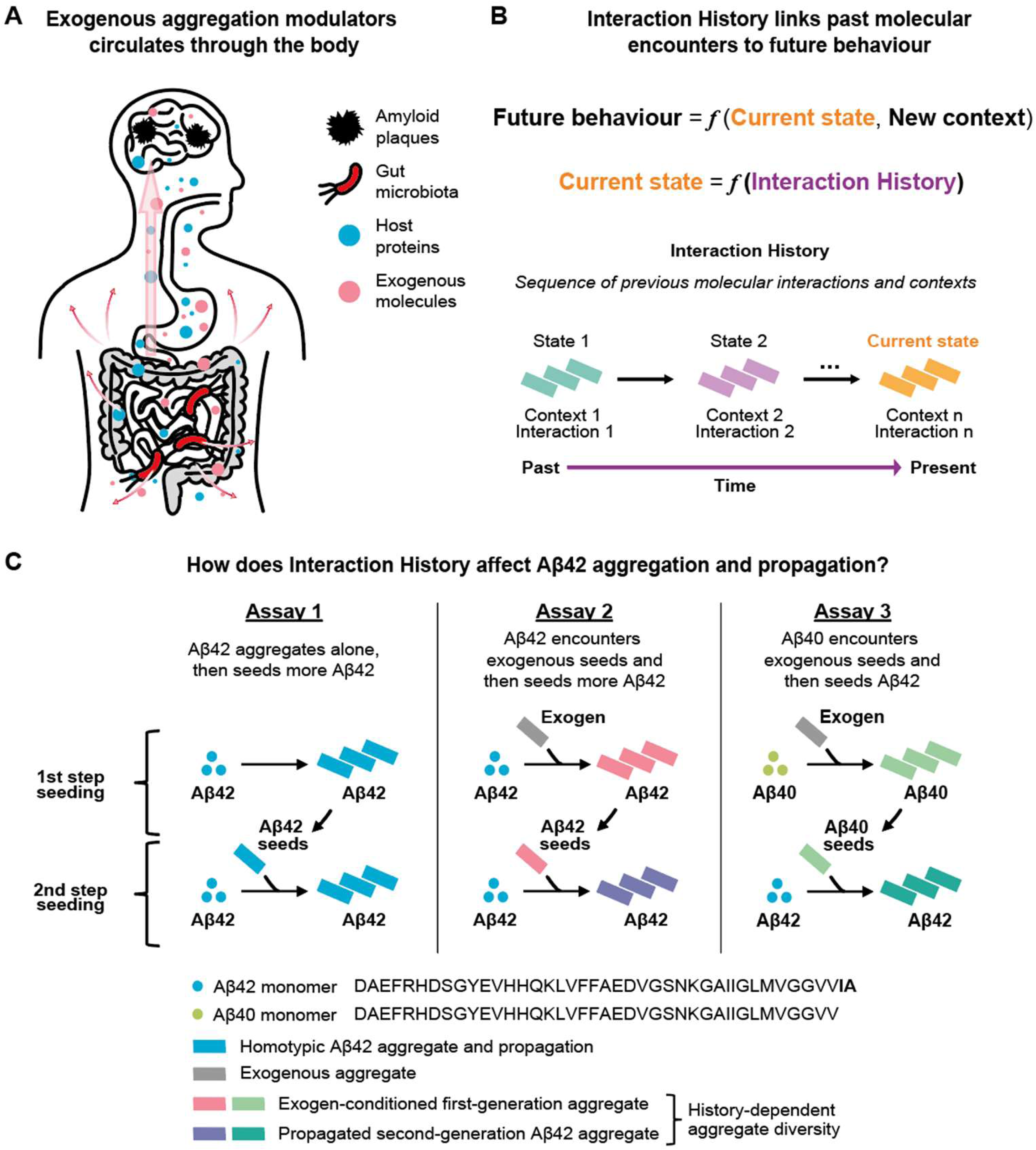
Interaction History in Aβ aggregation and propagation. Conceptual overview of the biological context (A), the principle of Interaction History (B), and the experimental design used to examine it (C). A) Microbial cells and their products can circulate through the body (pink arrows), encounter host proteins, reach distant tissues including the brain, and potentially influence protein aggregation processes. B) Conceptual representation of Interaction History as the dependence of the current molecular state on previous interactions and contexts. The future behavior of an aggregate depends on its current state and the new molecular context it encounters, while the current state itself reflects the sequence of molecular interactions experienced during earlier aggregation events. The schematic illustrates this history as a progression through successive states and contexts over time, culminating in the present aggregate state. C) Schematic representation of the sequential seeding approach used to examine how Interaction History influences Aβ42 aggregation and propagation. The sequences of Aβ40 and Aβ42 are shown, highlighting the two additional C-terminal residues of Aβ42. Three aggregation histories are considered. Assay 1: Aβ42 aggregates in the absence of exogenous peptides and subsequently undergoes canonical homotypic propagation. Assay 2: Aβ42 aggregates in the presence of an exogenous peptide, generating peptide-conditioned first-generation aggregates that subsequently seed monomeric Aβ42. Assay 3: Aβ40 aggregates in the presence of an exogenous peptide, generating peptide-conditioned first-generation aggregates that subsequently seed Aβ42. Although the second-generation reaction contains the same Aβ42 monomer substrate in all three assays, differences generated during the first aggregation step may influence the properties of the propagated Aβ42 assemblies, resulting in history-dependent aggregate diversity. Dots represent Aβ monomers and bars represent aggregated species. Colors distinguish Aβ isoforms, exogenous aggregates, peptide-conditioned first-generation aggregates, and the corresponding propagated second-generation Aβ42 aggregates.

Aβ aggregation is also influenced by interactions with other molecules, including proteins like amylin [16] and α-synuclein [17–19], nucleic acids [20,21], metals [22–24], and physicochemical parameters such as pH and ionic strength [25–29]. These factors can promote the formation of fibril polymorphs with distinct structural and kinetic profiles. Although *in vivo* systems tend to exhibit lower polymorphic variability, structurally distinct amyloid strains have been associated with divergent clinical manifestations in AD [10,30]. Emerging evidence suggests that exogenous molecules, including those derived from the microbiome, may modulate Aβ aggregation [31–33]. Microbial proteins with prion-like or amyloidogenic properties can become accessible within the intestinal environment through active secretion or following bacterial cell lysis [34,35]. Moreover, these proteins may be proteolytically processed into peptides capable of disseminating beyond the intestinal environment [36,37]. In the context of the gut–brain axis, microbiota-derived molecules may therefore encounter host proteins in molecular environments distinct from those in which they originated and, in some cases, reach distant tissues including the central nervous system (Figure 1A-B) [32,33,38–42]. In particular, microbiota-derived peptides with prion-like sequence features, such as low-complexity regions or aggregation-prone cores [32,39], represent potential modulators of protein aggregation. These sequences may act as heterotypic modulators or participate in cross-seeding events, in which preformed aggregates of the exogenous proteins can alter or trigger the aggregation of host amyloid-forming proteins such as Aβ (Figure 1C). While the biological significance of these interactions is still not fully understood, their effects on aggregation can be systematically studied under controlled *in vitro* conditions.

Amyloid assembly is not linear (Figure 2A) but rather dynamic and iterative [16–24,28,29], shaped by an energy landscape containing multiple accessible conformational states [28,29,43]. Molecular interactors and environmental changes can perturb this landscape by stabilizing particular intermediates or redirecting aggregation pathways. In biological systems, these perturbations can occur across different temporal and molecular contexts, as aggregates may encounter new molecular partners and act as seeds in later aggregation events. When such changes influence successive aggregation cycles, the history of previous molecular interactions becomes relevant to the subsequent reactions. We define this dependence as “Interaction History”: the influence of molecular interactions experienced during earlier aggregation cycles on the properties of subsequently propagated aggregates (Figure 1B). Mechanistically, prior interactions may alter the distribution of conformations generated in one cycle, thereby changing the properties of the seeds transmitted to the next.

**Figure 2.**
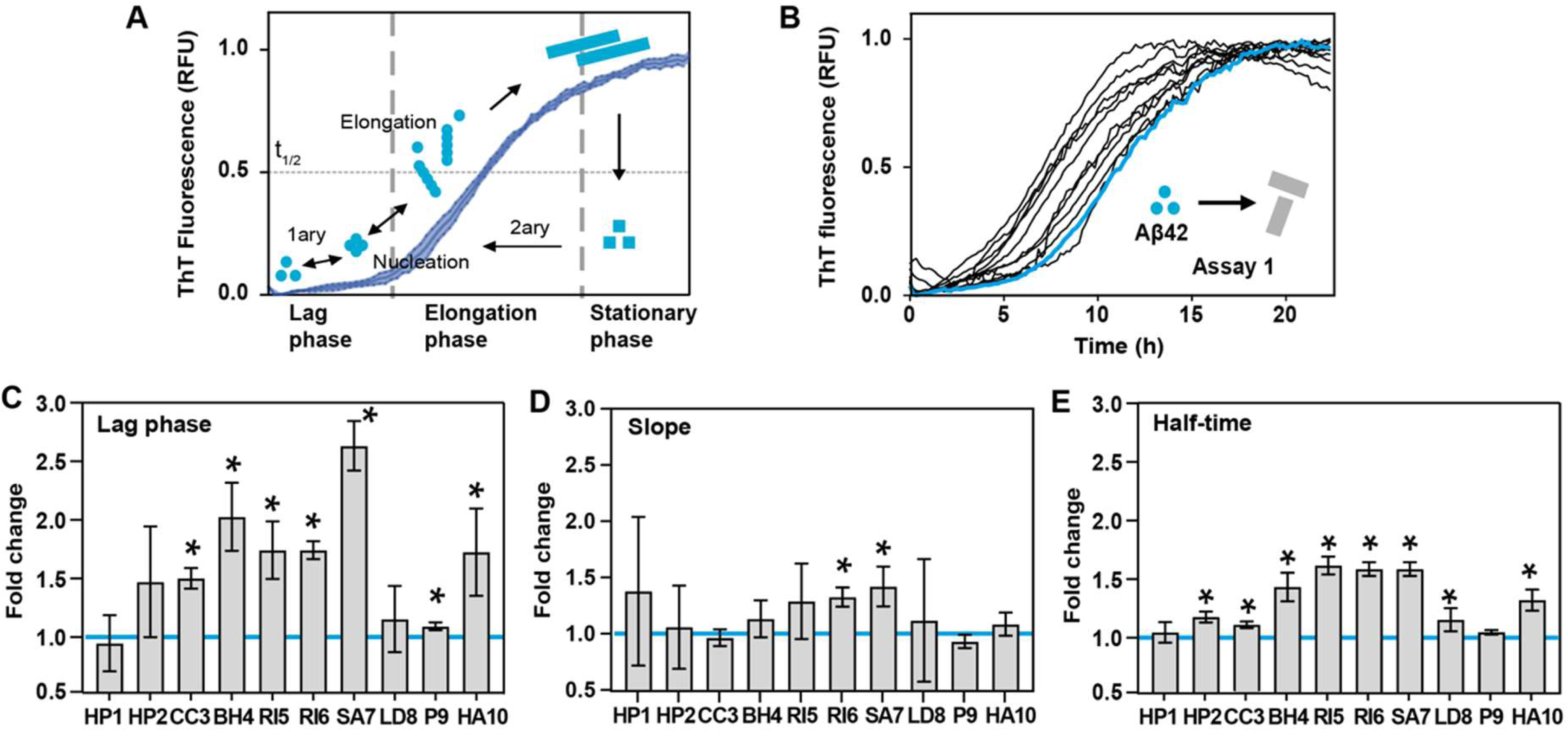
Exogenous Prion-like Sequences Influence Aβ42 Aggregation. A) Diagram showing the different phases of a protein aggregation process, indicating the different assembly mechanisms involved. B) Aggregation kinetics of Aβ42 in the presence (HP1–HA10, black lines) and absence (w/o, blue line) of seeds from exogenous prion-like sequences. See Supplementary Figure S4 for individual curves. C) Fold change in lag time (unseeded/seeded). D) Fold change in slope (seeded/unseeded). The equation fittings are shown in Supplementary Figure S5. E) Fold change in half-time (unseeded/seeded). The blue line serves as a visual reference indicating a fold change of one relative to the aggregation assay started only with Aβ42 monomers (unseeded). Each condition represents three biological replicates, with three technical replicates per biological replicate. Statistical comparisons were performed using multiple unpaired two-tailed t-tests with Benjamini–Krieger–Yekutieli FDR correction (Q = 0.05). An asterisk (*) indicates a significant difference relative to unseeded Aβ42. Kinetic parameters were expressed as fold changes relative to unseeded controls: slope was calculated as seeded/unseeded, whereas lag time and half-time were calculated as unseeded/seeded. With this convention, higher values for all three kinetic parameters indicate aggregation acceleration.

In our previous investigation of the gut microbiome, we built a collection of ten exogenous prion-like sequences derived from amyloidogenic regions of bacterial proteins [39]. These sequences were selected based on the amyloid stretch hypothesis, which states that specific regions within a protein, known as amyloid cores, can drive amyloid nucleation [44–46]. Using this collection, we showed that several bacterial peptides accelerated Aβ40 aggregation and impaired memory in *C. elegans* [39]. In the present study, we use this collection to test whether the molecular context in which an aggregate is initially formed influences its behavior when propagated under the same conditions. Using sequential heterotypic seeding assays, we examine how differences introduced during the first aggregation cycle influence the subsequent propagation in a common Aβ42 substrate, revealing differences in aggregation kinetics, conformational properties, and toxic potential (Figure 1C; Figures 2–7). By integrating these measurements across successive aggregation cycles, our findings suggest that single-step assays may miss aggregate diversity that becomes apparent only through successive reactions.

**Figure 3.**
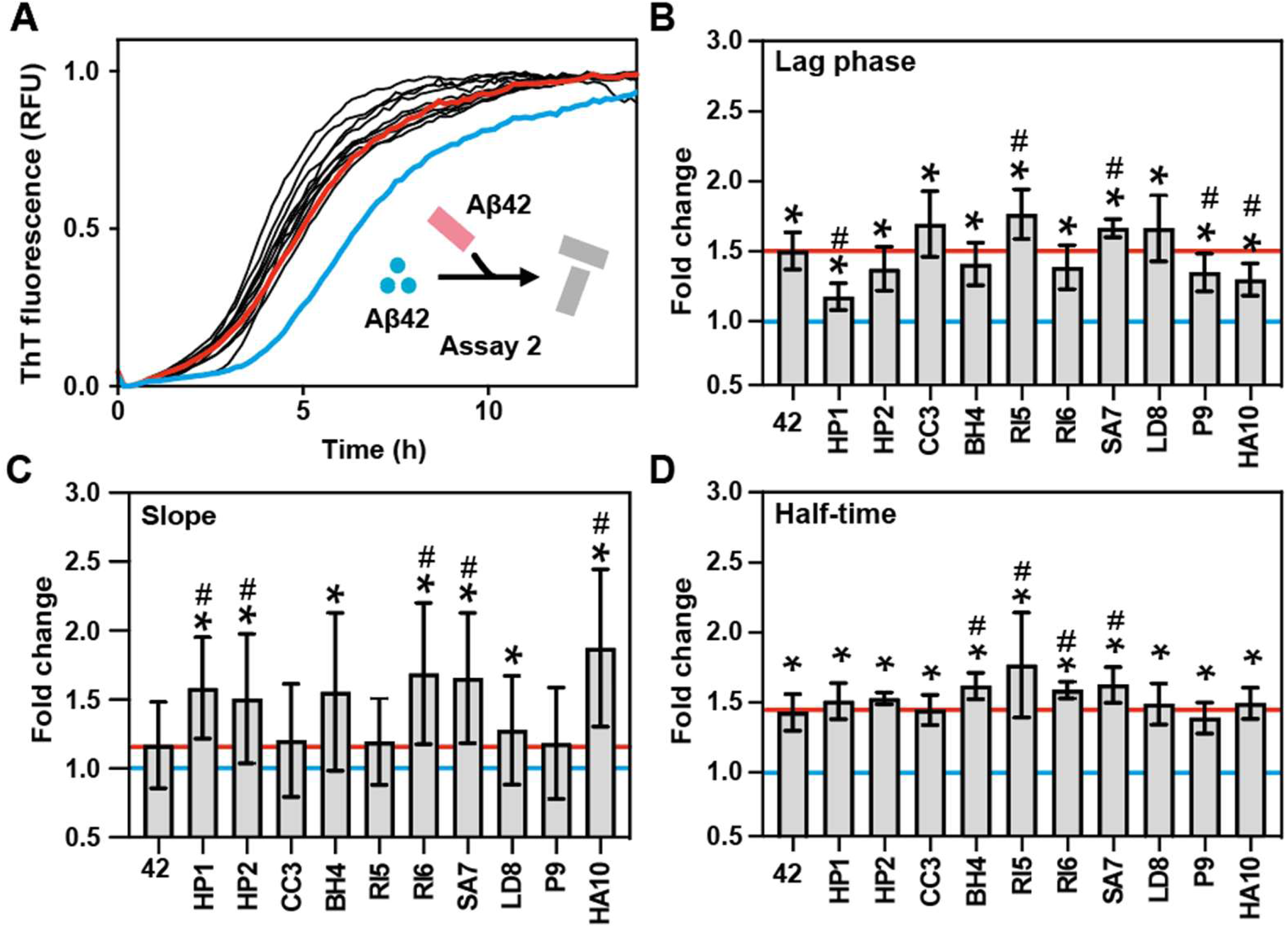
Exogenous Prion-like Sequences Affect Aβ42 Seeds and Their Propagation. A) Aggregation kinetics of Aβ42 seeded with Aβ42 aggregates formed in the presence and absence (w/o) of seeds from exogenous prion-like sequences. In blue and red are highlighted the aggregation kinetics of Aβ42 without seeds (negative control) and with seed of pure Aβ42 aggregates (homotypic seeding control), respectively. See Supplementary Figure S6 for individual curves. B) Fold change in lag time (unseeded/seeded). C) Fold change in slope (seeded/unseeded). The equation fittings are shown in Supplementary Figure S7. D) Fold change in half-time (unseeded/seeded). The horizontal blue line indicates a one-fold change, corresponding to unseeded Aβ42 aggregation. The horizontal red line indicates the mean value of the Aβ42 homotypic seeding control (42), corresponding to Aβ42 seeded with pure Aβ42 aggregates. Error bars indicate standard deviation. Each condition represents three biological replicates, with three technical replicates per biological replicate. Statistical comparisons were performed using multiple unpaired two-tailed t-tests with Benjamini–Krieger–Yekutieli FDR correction (Q = 0.05). An asterisk (*) indicates a significant difference relative to unseeded Aβ42, whereas a hash symbol (#) indicates a significant difference relative to the Aβ42 homotypic seeding control. Kinetic parameters were expressed as fold changes relative to unseeded controls: slope was calculated as seeded/unseeded, whereas lag phase and half-time were calculated as unseeded/seeded. With this convention, higher values for all three kinetic parameters indicate aggregation acceleration.

**Figure 4.**
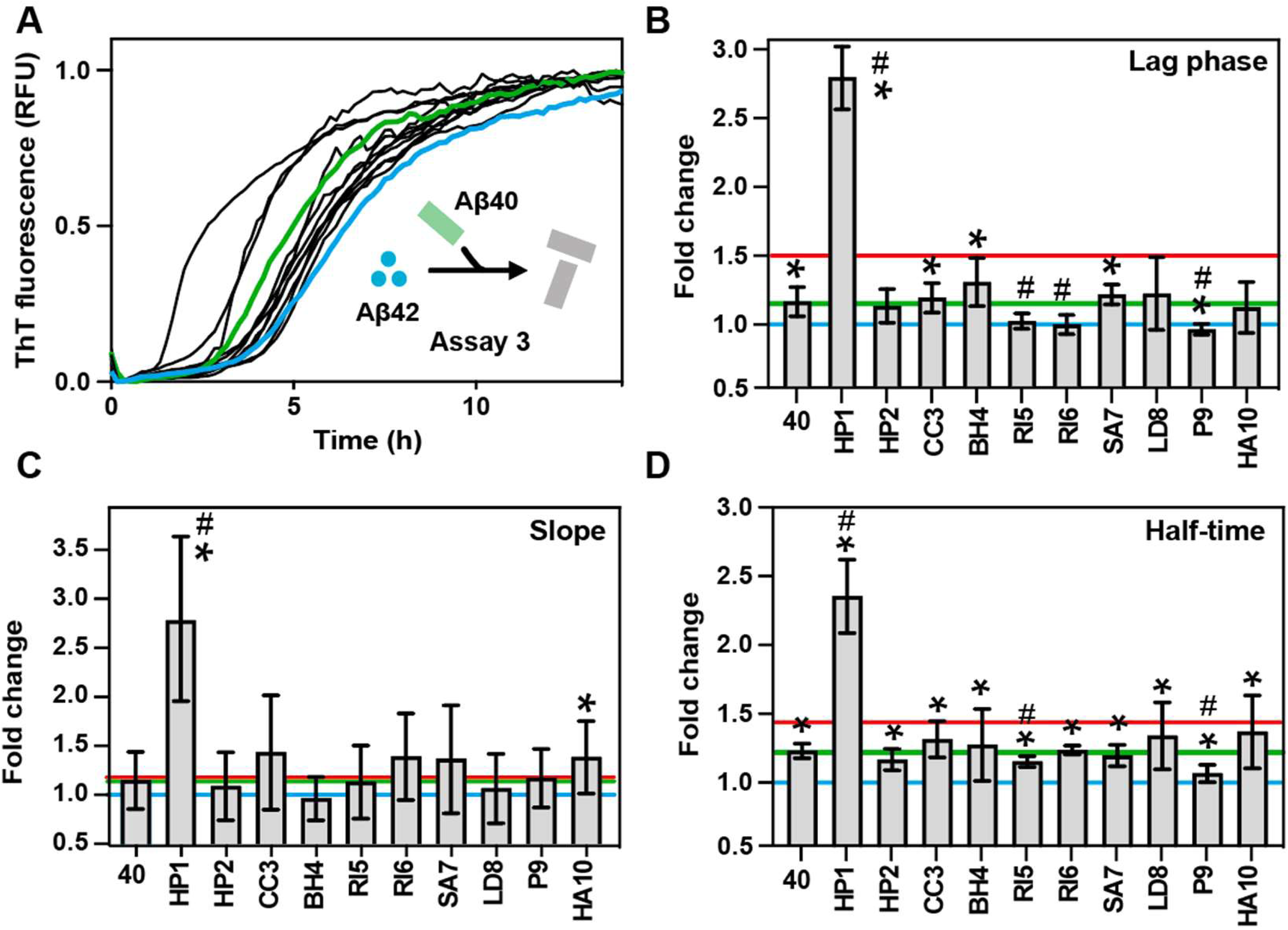
Exogenous Prion-like Sequences Affect Aβ40 Aggregation and Its Propagation to Aβ42. A) Aggregation kinetics of Aβ42 seeded with Aβ40 aggregates formed in the presence (HP1–HA10) and absence (w/o) of seeds from exogenous prion-like sequences. See Supplementary Figure S8 for individual curves. Green and blue lines highlight the aggregation of Aβ42 seeded with pure Aβ40 aggregates or without seeds, respectively. B) Fold change in lag time (unseeded/seeded). C) Fold change in slope (seeded/unseeded). The equation fittings are shown in Supplementary Figure S9. D) Fold change in half-time (unseeded/seeded). The horizontal lines indicate the average value for aggregation kinetics of Aβ42 seeded with pure Aβ42 aggregates (red, homotypic seeding), pure Aβ40 aggregates (green) or without seeds (blue, negative control). Error bars indicate the standard deviation. Each condition represents the results of three biological replicates, with three technical replicates per biological replicate. Statistical comparisons were performed using multiple unpaired two-tailed t-tests with Benjamini–Krieger–Yekutieli FDR correction (Q = 0.05). An asterisk (*) indicates a significant difference relative to unseeded Aβ42, whereas a hash symbol (#) indicates a significant difference relative to the Aβ42 homotypic seeding control. Kinetic parameters were expressed as fold changes relative to unseeded controls: slope was calculated as seeded/unseeded, whereas lag time and half-time were calculated as unseeded/seeded. With this convention, higher values for all three kinetic parameters indicate aggregation acceleration.

**Figure 5.**
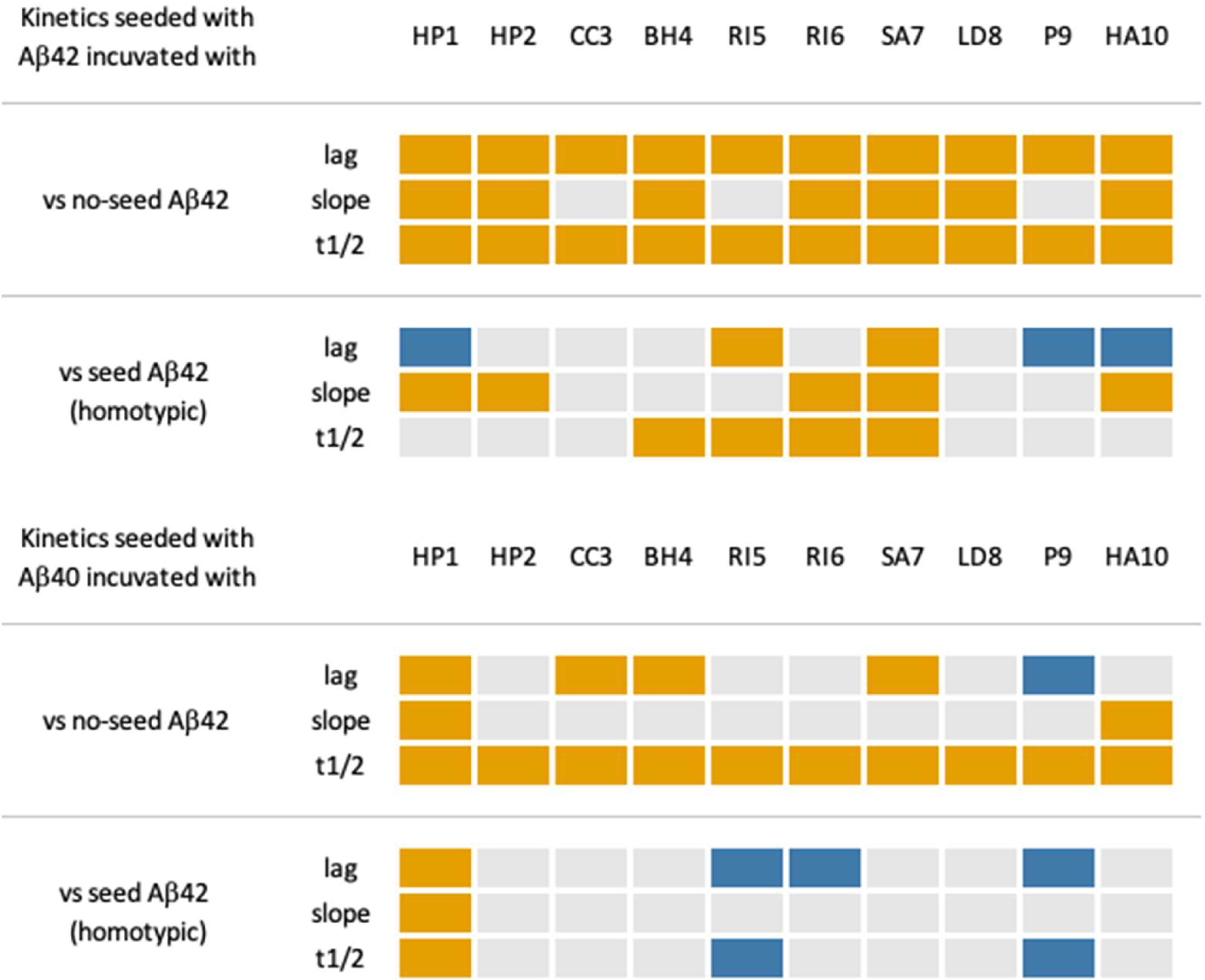
Differential Effects of Exogenous Prion-Like Sequences on Aβ42 Aggregation. Heat maps summarizing the statistically significant changes in the kinetic parameters measured during the second Aβ42 aggregation reaction (Figures 3 and 4). The upper panel corresponds to second-step Aβ42 reactions seeded with first-generation Aβ42 aggregates generated in the presence of the indicated bacterial peptides. The lower panel corresponds to second-step Aβ42 reactions seeded with peptide-conditioned Aβ40 aggregates. Columns indicate the bacterial peptide present during the first aggregation step. Each panel includes two complementary comparisons. “vs no-seed Aβ42” compares the aggregation reaction with unseeded Aβ42 (negative control) and evaluates whether first-generation aggregates influence Aβ42 aggregation. “vs seed Aβ42” compares the same reactions with Aβ42 seeded by homotypic Aβ42 aggregates (homotypic seeding control) and evaluates how bacterial peptides modify the propagation properties of the resulting aggregates relative to canonical Aβ42 seeds. Rows correspond to the three kinetic parameters analyzed (lag phase, slope, and half-time). Orange indicates a statistically significant acceleration of Aβ42 aggregation, blue indicates a statistically significant deceleration, and white indicates no significant difference. For lag phase and half-time, acceleration corresponds to reduced time values, whereas for slope it corresponds to an increased slope.

**Figure 6.**
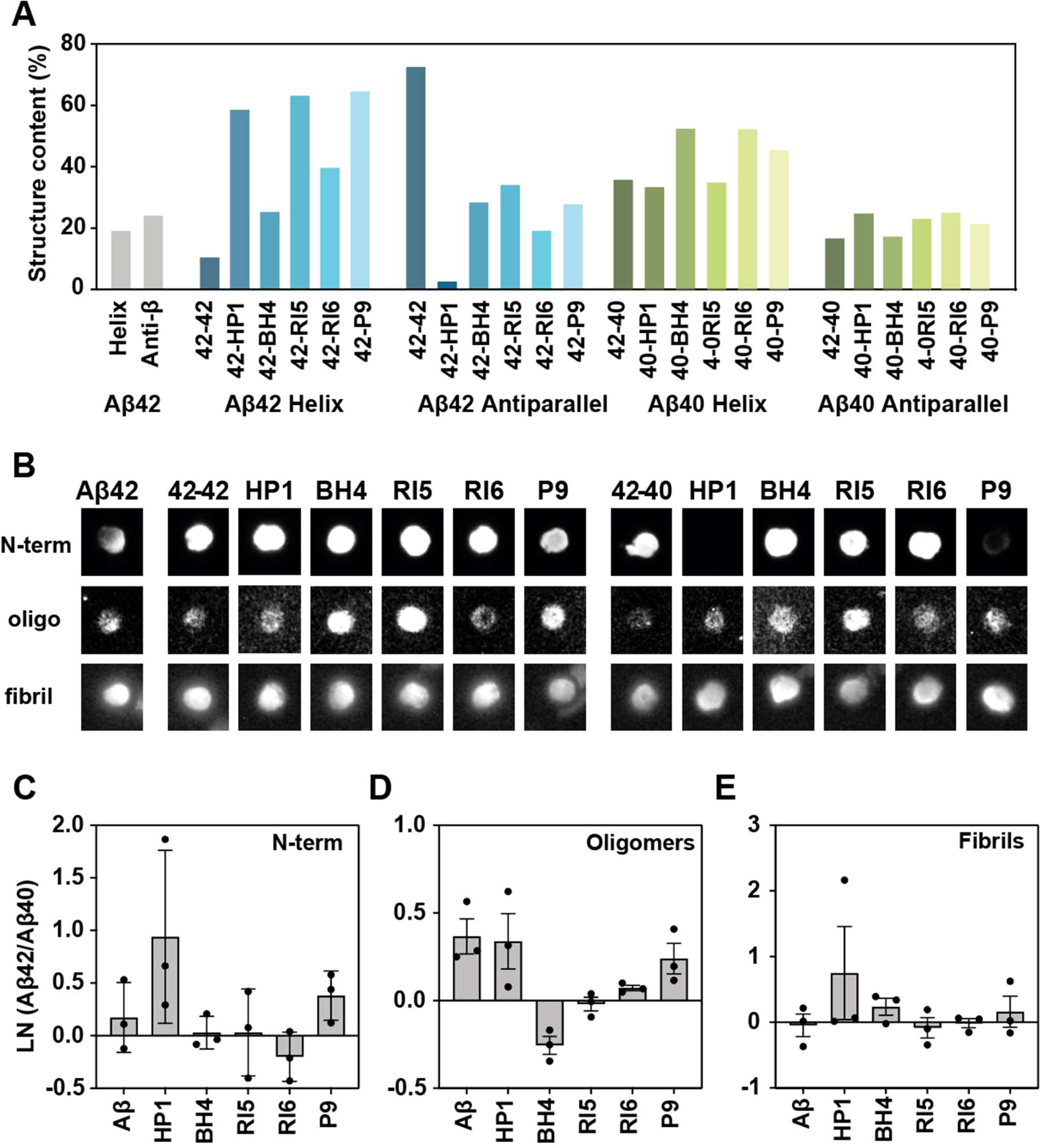
Secondary-structure content and epitope accessibility of propagated Aβ42 aggregates. (A) Circular dichroism (CD) spectral deconvolution of end-point material from each aggregation assay. Bars indicate the estimated contributions of helix-like and antiparallel β-like structures. (B) Representative dot blots of end-point material probed with three conformation/epitope-sensitive antibodies: anti-N-terminal (6E10, epitope 3–8; “N-term”), anti-oligomer (A11; “oligo”), and anti-fibril (anti-amyloid fibrils antibody; “fibril”). Conditions correspond to second-generation Aβ42 assemblies seeded with first-generation aggregates generated from either Aβ42 (left block; 42-42 and 42–peptide) or Aβ40 (right block; 42-40 and 40–peptide) in the presence of the indicated exogenous peptides (HP1, BH4, RI5, RI6, and P9). (C–E) To directly compare Aβ42- and Aβ40-derived seeding histories for each peptide, matched dot-blot signals within each independent experiment were expressed as ln(Aβ42/Aβ40). This transformation centers equivalent reactivity at 0 and provides a symmetric representation of reciprocal changes: values >0 indicate higher signal in the Aβ42-derived condition, whereas values <0 indicate higher signal in the Aβ40-derived condition. Panels show N-terminal (C), oligomer (D), and fibril (E) reactivity. Bars represent the mean across three independent experiments, points indicate individual experiments (n = 3), and error bars indicate SD.

**Figure 7.**
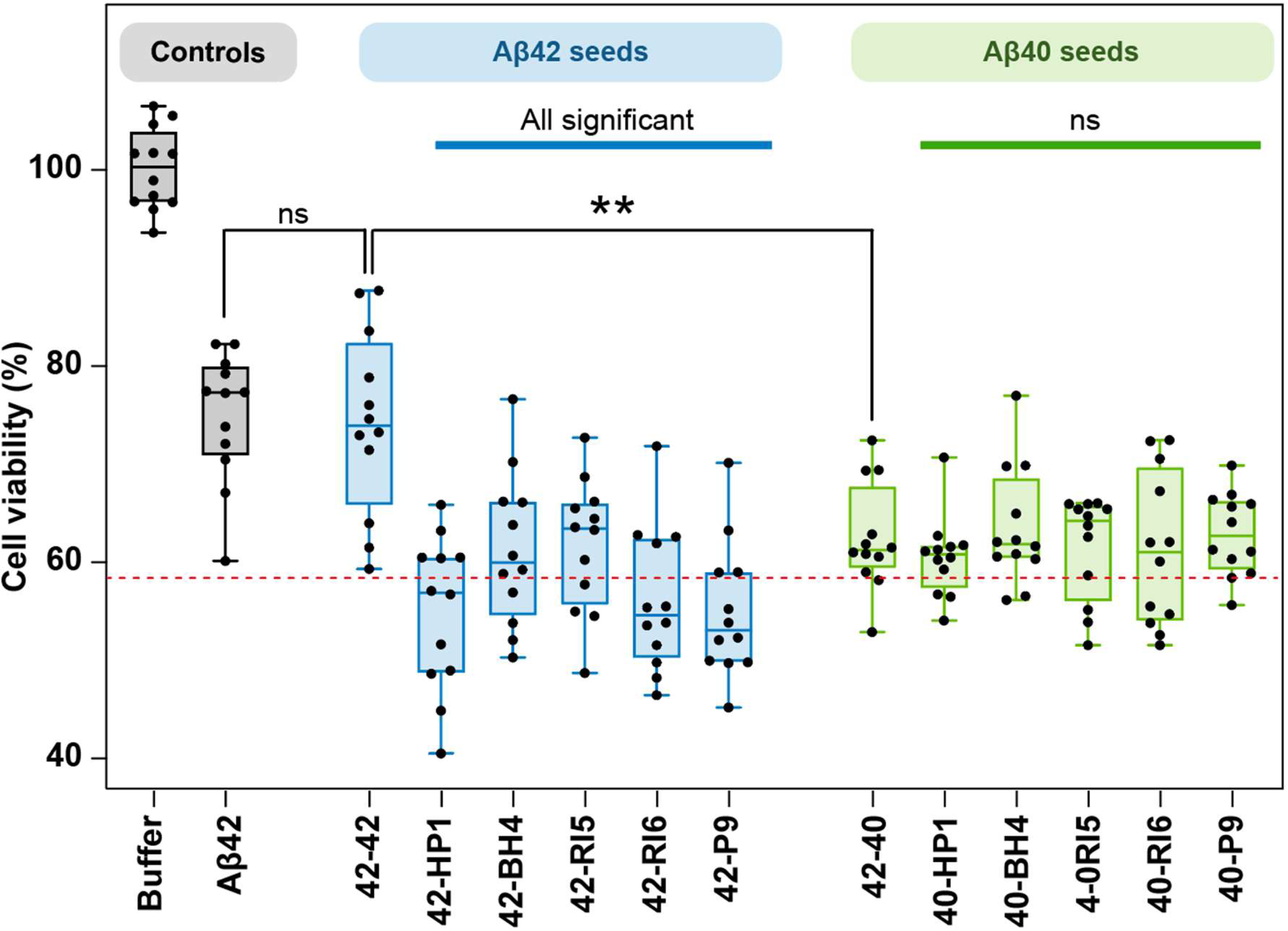
Toxicity of peptide-conditioned Aβ42 aggregates in neuron-like SH-SY5Y cells. Neuron-like differentiated SH-SY5Y neuroblastoma cells were exposed for 24 h to end-point material from the second aggregation reaction seeded with first-generation aggregates generated from either Aβ42 (blue; Aβ42 seeds) or Aβ40 (green; Aβ40 seeds) in the presence of the indicated bacterial peptides (HP1/C1, BH4/C4, RI5/C5, RI6/C6, and P9/C9). Buffer, unseeded Aβ42, and homotypic Aβ42-seeded conditions are shown as controls (grey). Cell viability was quantified by MTT and expressed as a percentage relative to the buffer-only condition. Box plots show the median and interquartile range; whiskers indicate the range, and individual points represent measurements from three independent experiments with four technical replicates each (n = 12). The red dashed line marks the overall mean viability across all aggregate-treated conditions and is included as a visual reference. Thick colored horizontal bars summarize the individual comparisons of each peptide-conditioned sample with the corresponding peptide-free seeding control. “All significant” indicates that all five Aβ42-derived peptide-conditioned samples differed significantly from the homotypic Aβ42-seeded control (42–42), whereas “ns” indicates that none of the Aβ40-derived peptide-conditioned samples differed significantly from the peptide-free Aβ40-derived control (42–40). The thin black lines indicate the comparison between the no-seeded and canonical-seeded conditions (left, not significant) and between the canonical-seeded conditions and the Aβ40 homotypic seeds (right, p < 0.01). Statistics were performed using ordinary one-way ANOVA followed by Dunnett’s multiple-comparisons test (ns, not significant; * p < 0.05; ** p < 0.01).

## 2. Results and Discussion

### 2.1. Exogenous Prion-like Sequences Can Influence Aβ Aggregation Kinetics

To further investigate how exogenous amyloidogenic sequences influence Aβ42 aggregation, we used the previously characterized collection of ten gut microbiota-derived prion-like peptides [39] (Table 1). The peptides comprise 21-residue amyloidogenic regions with net charges ranging from +2 to −2 and compositional similarities below 30% (Table 1), providing a controlled set of chemically diverse exogenous sequences [39].

**Table 1.**
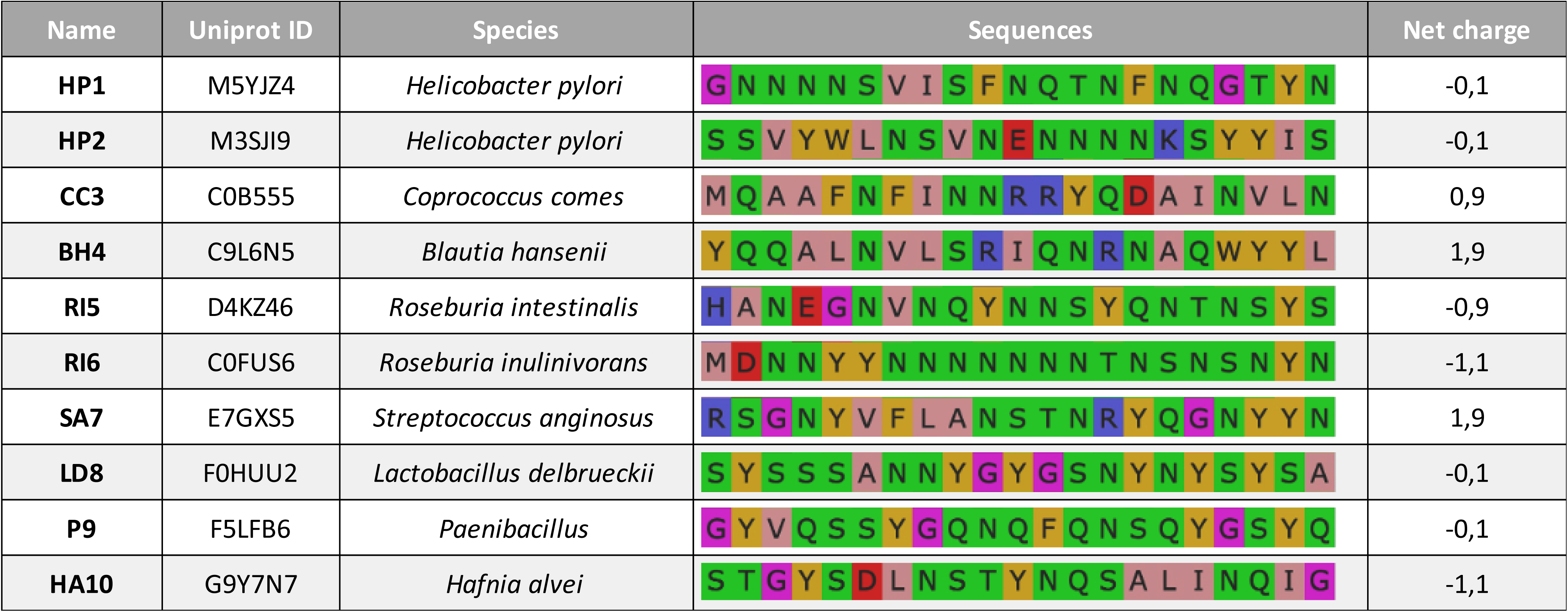
Set of Exogenous Prion-like Sequences. Columns (from left to right) display: sequence identifiers used in this study; UniProt accession codes; source species from which the sequences originate; amino acid sequences; and net charge measured with Protein Calculator V3.4. The amino acids are colored according to the Zappo color scheme: aliphatic/hydrophobic (ILVAM) - rose, aromatic (FWY) - orange, positive (KRH) - dark blue, negative (DE) - red, hydrophilic (STNQ) - light green, conformationally special (PG) - magenta, cysteine (C) - yellow. The sequences were obtained from Curto, J.S. et al. [39].

To model the sequential and context-dependent nature of aggregation in biological systems, we used heterotypic seeding assays that generate seeds through different prior molecular interactions and subsequently propagate them in a common Aβ42 substrate (Figures 1B and 1C). We incorporated the interplay between Aβ42 and Aβ40, hypothesizing that the higher abundance of Aβ40 [4,6] may increase its probability of encountering exogenous molecules and thereby indirectly influence subsequent Aβ42 aggregation. To explore this interplay, we monitored Thioflavin-T (ThT) fluorescence across successive aggregation reactions (Figures 2– 4) and used transmission electron microscopy (TEM) to examine the resulting aggregates (Supplementary Figures S1 and S2).

In our previous study [39], we found that most of the bacterial prion-like sequences (eight out of ten) accelerated Aβ40 aggregation, with lag time correlating with the sequences’ net charge. Hence, positively charged sequences led to faster aggregation, while negatively charged ones had little effect or even slowed aggregation [47–49]. These results highlight the role of electrostatic forces in intermolecular interactions and align with earlier reports showing that negatively charged peptides inhibit Aβ aggregation [47,48]. In the present study, we extended our analysis to Aβ42 by incubating it with these pre-aggregated bacterial peptides (Figure 2B). As observed with Aβ40, incubation of Aβ42 with a 1:10 molar ratio of prion-like sequences predominantly enhances its aggregation (eight samples presented shorter half-time, Figure 2E).

Unlike the experiments performed with Aβ40 [39], where aggregation kinetics correlated with the peptide net charge, the association in Aβ42 was weaker and was detected only when considering absolute net charge (Supplementary Figure S3). Nevertheless, the two most positively charged peptides, SA7 and BH4 [39], accelerated Aβ42 aggregation, producing the shortest lag phases and low half-times (Figure 2C–E). Overall, these results suggest that electrostatic properties contribute to the modulation of Aβ aggregation but are not sufficient to explain the full range of peptide-specific effects.

From a global point of view, the acceleration observed in Aβ42 aggregation was mainly driven by shortening the lag phase: seven samples showed reduced lag times, whereas only two displayed steeper slopes (Figure 2B–D). A reduced lag time indicates a more efficient primary nucleation phase, involving oligomeric species that serve as templates, whereas a steeper slope indicates a faster aggregation rate during the growth phase, which may result from the rapid addition of monomers to existing fibrils and could also involve secondary nucleation events (Figure 2A) [50,51]. The half-time (t1/2) serves as an indicator of overall aggregation efficiency, influenced by both lag phase and slope. Together, these findings suggest that these exogenous prion-like sequences primarily enhance the nucleation of Aβ42 aggregation by acting as seeding points (Figure 2).

### 2.2. Effect on Aβ42 Seeds and Their Propagation

After observing that bacterial prion-like sequences can directly influence the aggregation of both Aβ40 [39] and Aβ42, we investigated how past molecular interactions affect the propagation of Aβ42 aggregates (Figure 1C). To address this, we conducted kinetic assays using pre-formed Aβ fibrils previously seeded with the bacterial peptides (Figure 2, Supplementary Figure S4). This second aggregation step (Figure 1C and Figure 3) involved a tenfold dilution of the exogenous peptides compared to the initial direct seeding assays. This dilution was achieved by incorporating 10% of the Aβ aggregates formed during the first step (Methods), minimizing the direct influence of the bacterial molecules (≈1%) and allowing us to focus on the properties of the pre-formed aggregates.

All lag times were accelerated compared with Aβ42 without seeds (Figure 3B, * symbol), with two bacterial sequences (RI5 and SA7) producing significantly shorter lag phases than the homotypic Aβ42-seeded reaction (Figure 3B, # symbol). Regarding the slope, seven sequences exhibited significantly steeper slopes than unseeded Aβ42, while five out of ten produced significantly steeper slopes than the homotypic Aβ42-seeded control. Together, these results show that peptide-conditioned Aβ42 aggregates can propagate altered seeding properties in the subsequent aggregation reaction, with some conditions producing greater aggregation acceleration than homotypic Aβ42 seeds. This also underscores the importance of the Interaction History, demonstrating that even exogenous molecules with limited initial seeding capacity can redirect the aggregation pathway toward aggregates with enhanced propagation potential. Additionally, whereas the first aggregation cycle (Figure 2), seeded directly with the bacterial peptides, predominantly affected the lag phase, the second cycle (Figure 3), seeded with the resulting Aβ42 aggregates, resulted in a more consistent acceleration across peptide conditions, affecting both lag phase and slope. This increased seeding efficiency is consistent with the greater sequence compatibility between the Aβ42-containing seeds and the Aβ42 monomer substrate [52–54].

Beyond this general increase in seeding efficiency, peptide-conditioned Aβ42 seeds displayed distinct kinetic profiles when compared with homotypic Aβ42 seeding (Figure 3A–D; Supplementary Figure S6). RI5 and SA7 shortened the lag phase, whereas HP1, P9, and HA10 prolonged it. In contrast, HP1, HP2, RI6, SA7, and HA10 increased the slope. Thus, individual peptide-conditioned seeds could affect different phases of the aggregation reaction, and even in opposite directions, as observed for HP1 and HA10, which prolonged the lag phase while increasing the slope. Changes in half-time reflected the combined kinetic outcome, with significant acceleration observed for BH4, RI5, RI6, and SA7. CC3 and LD8 showed no significant differences from homotypic Aβ42 seeding. These results indicate that exposure to different exogenous peptides does not uniformly enhance or reduce Aβ42 seeding efficiency, but generates distinct kinetic profiles during subsequent propagation. Importantly, these differences remain detectable in the second aggregation reaction, in which the exogenous peptides are substantially diluted. The differential effects on lag phase and slope further indicate that peptide-conditioned seeds can influence distinct stages of subsequent Aβ42 aggregation. Such history-dependent kinetic differences may contribute to the diversity of the resulting Aβ42 assemblies.

### 2.3. Peptide-conditioned Aâ40 aggregates show distinct effects on subsequent Aâ42 aggregation

While the above experiments focused on the self-seeding of Aβ42, it is important to consider that Aβ40 is the predominant isoform of amyloid-beta peptide in the human body [4,6]. Given its higher abundance, Aβ40 is more likely to encounter and interact with exogenous prion-like proteins and may therefore represent an additional route through which exogenous molecules influence subsequent Aβ42 aggregation. To explore this possibility, we examined whether our collection of exogenous sequences could also influence the cross-seeding potential of Aβ40 on Aβ42 (Figure 4, Supplementary Figure S8).

In our previous work, we observed that most bacterial sequences from our collection accelerated the aggregation of monomeric Aβ40 [39]. Here, we used Aβ40 aggregates formed during this initial heterotypic aggregation reaction to seed Aβ42 deposition. As a control, we tested the seeding efficiency of homotypic Aβ40 aggregates and found that, although they reduced the lag time and half-time of Aβ42 aggregation to a lesser extent than Aβ42 seeds (Figure 4B and D), they induced a comparable slope increase (Figure 4C). These findings support that the two additional C-terminal residues of Aβ42 play a critical role in facilitating the assembly of free monomers during the primary nucleation process, and their absence in Aβ40 reduces its propagation efficiency [55].

Peptide-conditioned Aβ40 seeds produced a markedly different kinetic pattern from the corresponding Aβ42-derived seeds. Relative to unseeded Aβ42, four conditions (HP1, CC3, BH4, and SA7) shortened the lag phase, whereas P9 prolonged it (Figure 4B). Only HP1 and HA10 significantly increased the slope (Figure 4C), while all ten peptide-conditioned Aβ40 seeds reduced the half-time of Aβ42 aggregation (Figure 4D). When compared directly with homotypic Aβ42 seeding, HP1 was the only Aβ40-derived condition that further accelerated all three kinetic readouts. In contrast, RI5 and P9 prolonged both lag phase and half-time, while RI6 selectively prolonged the lag phase (Figure 4, Supplementary Figure S8).

Overall, the summary of kinetic changes (Figure 5) reveals distinct propagation patterns depending on the first-generation seeding history. Peptide-conditioned Aβ42 seeds produced a broader range of kinetic responses, including both acceleration and deceleration across different parameters, whereas Aβ40-derived seeds showed a more restricted pattern of changes. Aβ40-derived seeds also showed generally lower propagation capacity, consistent with a putative regulatory or protective role for Aβ40 [9,12,13]. HP1 was the main exception, producing Aβ40-derived seeds that exceeded even homotypic Aβ42 seeding in several kinetic parameters. Interestingly, HP1 derives from a putative vacuolating cytotoxin sequence of *Helicobacter pylori* (Table 1), a bacterium previously linked to AD [56,57]. Consistent with this observation, our previous work showed that HP1 produced one of the strongest negative effects on cognitive performance in *C. elegans* [39].

### 2.4. Prion-Like Peptides Imprint Distinct Structural Signatures on Propagated Aβ Aggregates

To determine whether initial exposure to bacterial prion-like peptides results not only in kinetic modulation but also in structural differences in the propagated aggregates, we analyzed end-point material obtained after the two-step aggregation assay. For this purpose, we used circular dichroism (CD) to examine structural variability across conditions, complemented by dot-blot analysis to assess conformational features and epitope accessibility.

To capture the diversity of kinetic responses observed in the sequential seeding assays (Figure 5), we selected five peptide conditions (HP1, BH4, RI5, RI6, and P9) representing distinct aggregation behaviors for subsequent structural and toxicity analyses. HP1 was included because it is the only peptide that generates Aβ40-derived seeds with seeding efficiency comparable to or exceeding that of Aβ42 homotypic seeds. At the opposite extreme, P9 produces minimal acceleration in both Aβ40- and Aβ42-seeded reactions. In addition, BH4, RI5, and RI6 were chosen because they induce distinct and non-overlapping kinetic alterations: BH4 and RI6 display similar effects in Aβ42-seeded kinetics but diverge markedly when Aβ40-derived seeds are used, whereas RI5 exerts comparatively mild effects and can even decelerate aggregation in some Aβ40-seeded assays.

The CD spectra indicated differences in secondary-structure content across conditions, with an increased contribution of helix-like signatures in peptide-conditioned reactions, particularly within the Aβ42-derived samples (Figure 6). In addition, Aβ42-derived samples showed greater structural variability across peptide conditions than Aβ40-derived samples, particularly for the β-like component (coefficient of variation, 55% vs 14%; Figure 6). Among these, HP1 produced the most pronounced change in the Aβ42-derived samples relative to the homotypic Aβ42 reaction (Figure 6).

To further characterize the propagated assemblies, we examined specific conformational features by dot-blot analysis using three antibodies reporting on distinct aggregate properties: (i) an anti-oligomer antibody (A11), which recognizes prefibrillar oligomeric assemblies; (ii) an anti-amyloid fibrils antibody, which recognizes fibrillar cross-β structure; and (iii) an anti-N-terminal anti-Aβ antibody (6E10; epitope 3–8), used here as a proxy for N-terminal segment accessibility.

To enable comparisons across independent blots performed on three experimental days, all chemiluminescence signals were normalized to the Aβ42 no-seed condition. Under this normalization, ratios were generally higher in the anti-N-terminal readout than in the anti-oligomer or anti-fibril readouts (Supplementary Figure S10 and S11), reflecting the low anti-N-terminal signal of the Aβ42 no-seed reference (Figure 6B). At the level of individual conditions, anti-N-terminal ratios were typically ≥1 across most peptide-conditioned samples (Supplementary Figure S11), whereas anti-oligomer and anti-fibril ratios generally clustered close to or below 1. Notably, the anti-fibril readout showed the clearest difference in dispersion between seeding histories: Aβ40-seeded conditions were relatively homogeneous (with a standard deviation of 0.06), whereas Aβ42-seeded conditions were markedly more heterogeneous (standard deviation of 0.26, 4-fold higher).

To compare seeding histories directly for each peptide, we computed ln(Aβ42/Aβ40) for each antibody (Figure 6C-E). For the anti-N-terminal readout (Figure 6C), most peptides showed positive values, indicating higher 6E10 reactivity in the Aβ42-seeded aggregates; HP1 displayed the largest positive shift, whereas RI6 showed a negative value, indicating higher reactivity in the Aβ40-seeded aggregates. For the anti-oligomer readout, most Aβ42-seeded aggregates showed higher chemiluminescence signals (Figure 6D). In contrast, BH4 showed a pronounced negative value, and RI5 remained near zero. For the anti-fibril readout (Figure 6E), most conditions showed ln(Aβ42/Aβ40) values close to zero, indicating comparable anti-fibril reactivity between Aβ42- and Aβ40-derived seeding histories. HP1 was the main exception, showing higher anti-fibril reactivity in the Aβ42-derived condition.

We also examined whether aggregate conformation relates to aggregation kinetics. Overall, we found no consistent linear relationships, except for a correlation between the anti-oligomer signal in Aβ42-seeded aggregates and lag-phase duration (Supplementary Figure S12). This indicates that, within the Aβ42-derived samples, oligomer-like reactivity is associated with differences in aggregation onset. The lack of additional consistent correlations indicates that aggregate variability cannot be explained by simple linear relationships.

Overall, the dot-blot and CD profiles indicate that both secondary-structure content and epitope accessibility in the final Aβ42 material vary across conditions, and that the resulting conformational signatures depend on the identity of the exogenous peptide and the first-generation seeding history (Aβ42- vs Aβ40-derived seeds).

### 2.5. Different seeding histories generate Aβ aggregates with distinct toxic potential

From a biological perspective, the key question is whether the diversity of aggregation outcomes translates into differences in the toxic potential of the final assemblies. With the limitations of an *in vitro* system in mind, we analyzed the effects of these assemblies on cell viability in neuron-like differentiated SH-SY5Y neuroblastoma cells.

Peptide-conditioned Aβ42-derived samples showed lower viability (Figure 7) and greater heterogeneity than the corresponding Aβ40-derived samples (coefficient of variation 5.9% vs 2.0%; 3.0-fold higher). Within the Aβ42-derived samples, HP1 and P9 were the most toxic conditions and both also showed delayed aggregation onset. However, a significant correlation between lag phase and cell viability was observed only after excluding HP1 and RI5, indicating that toxicity is not explained by lag phase alone and likely depends on additional aggregate properties (Supplementary Figure S13 top). Within the Aβ40-derived samples, no correlation was observed between kinetic properties and cell viability, consistent with the fact that, despite their divergent kinetic profiles, these samples produced similar viability outcomes (Figure 7).

We next examined the relationship between aggregate conformation and toxicity. Anti-N-terminal signal positively correlated with cell viability both across all samples and within the Aβ42-derived subset (Supplementary Figure S14), suggesting that conditions with higher N-terminal accessibility are associated with reduced toxic potency. This aligns with previous studies showing that N-terminal features can influence Aβ aggregation and toxicity, through both local conformational presentation (epitope accessibility and surface properties) and sequence-level variation (N-terminal truncations or modifications) [27,58–60].

Within the Aβ42-derived samples, higher anti-oligomer signal was associated with lower cell viability (Supplementary Figure S13 bottom). Together with the relationship between lag-phase duration and anti-oligomer reactivity described above (Supplementary Figure S12), these observations are consistent with a model in which conditions that alter aggregation onset may favour a higher abundance of oligomer-like conformations with increased toxic potential [61–64]. In this context, both the identity of the exogenous peptide and the first-generation seeding history (Aβ40 vs Aβ42) appear to influence the abundance of these species.

### 2.6. Interaction History Shapes Aβ42 Properties

To obtain a global view of the results across the different seeding assays, we integrated kinetic parameters, epitope exposure, oligomer signal, and toxicity into an unsupervised principal component analysis (PCA). The PCA separated Aβ40-seeded from Aβ42-seeded samples and identified the two HP1-conditioned samples as distinct outliers (Figure 8). Notably, the homotypic Aβ42 control also occupied an extreme region of the PCA space.

**Figure 8.**
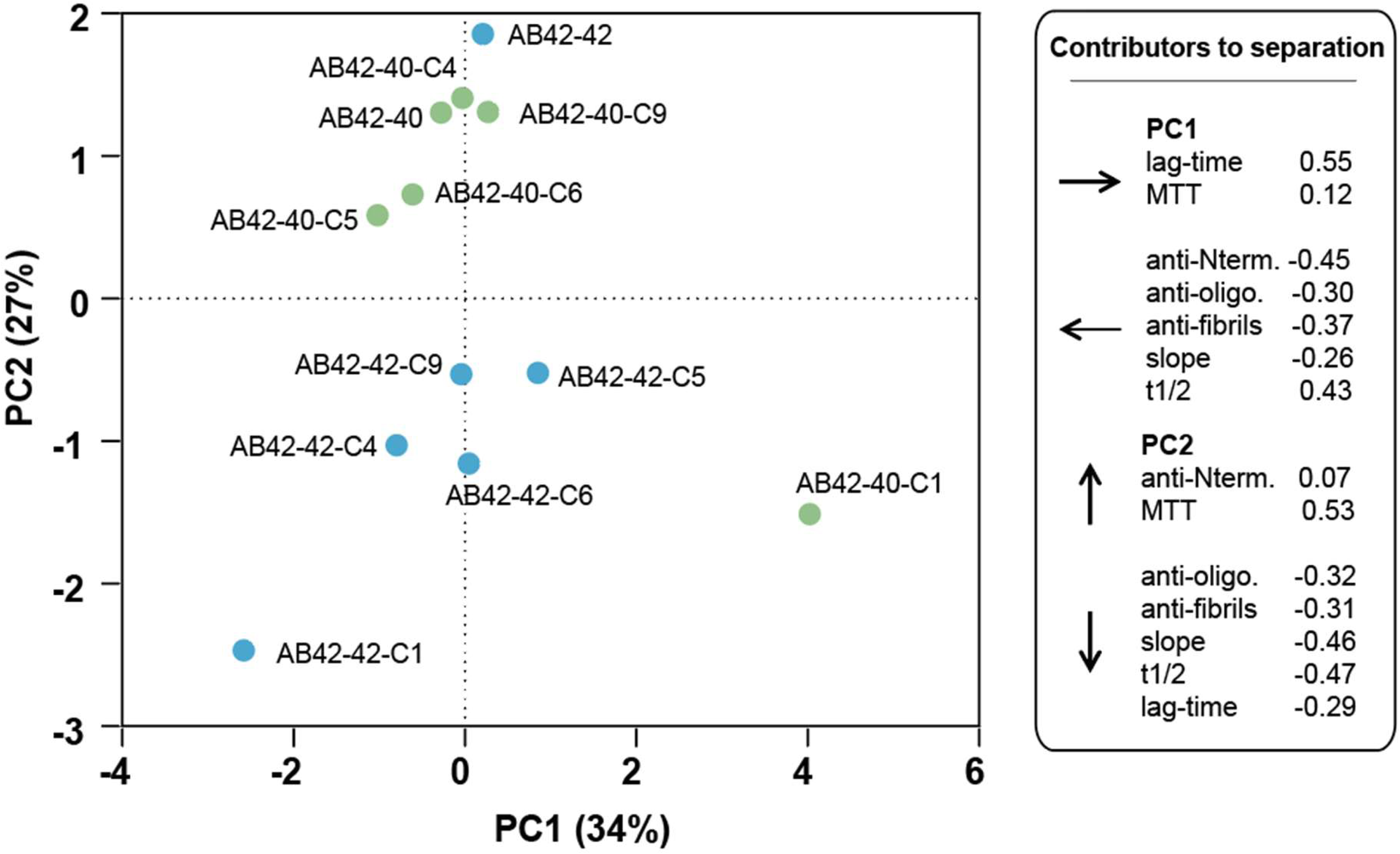
PCA integrating kinetic, epitope exposure, and toxicity features clusters Aβ42 assemblies derived from different seeding assays. Principal component analysis (PCA) was performed on variables derived from the two-step aggregation assay, including kinetic fold-changes (lag time, half-time, slope), dot-blot readouts (anti-N-terminal/6E10, anti-oligomer/A11, anti-fibril), and SH-SY5Y viability (MTT). Each point represents the end-point material from the second reaction (Aβ42 monomer) seeded with first-generation aggregates generated from either Aβ42 (blue) or Aβ40 (green) in the presence of the indicated bacterial peptide (C1/HP1, C4/BH4, C5/RI5, C6/RI6, C9/P9) or without peptide (AB42-42; AB42-40). PC1 and PC2 explain 34.4% and 27.3% of the variance, respectively. Variable loadings for each parameter are shown on the plot.

This separation was captured by the first two principal components, which together explained 61.7% of the variance (PC1, 34.4%; PC2, 27.3%). Accordingly, PC1 was driven primarily by the kinetic fold-changes in lag and half-time (positive loadings), opposed by the structural antibody readouts, most strongly the anti-N-terminal signal and, to a lesser extent, the anti-fibril and anti-oligomer signals (negative loadings). In contrast, PC2 was dominated by MTT viability (positive loading) and was opposed by slope and half-time fold-changes together with anti-oligomer and anti-fibril reactivity (negative loadings). Notably, the N-terminal signal contributed minimally to PC2 (Figure 8).

When projected onto the PC1–PC2 space, samples were distributed primarily along PC2. Aβ40 seeding assays clustered at higher PC2 values, indicating higher viability together with lower slope fold-change and reduced anti-oligomer and anti-fibril reactivity. By contrast, most Aβ42-derived conditions occupied lower PC2 values, corresponding to lower viability and comparatively higher slope fold-change, anti-oligomer reactivity, and anti-fibril reactivity. Variation along PC1 highlighted HP1-conditioned samples as extremes, with Aβ42-40-HP1 scoring highest on PC1 and Aβ42-42-HP1 scoring lowest, highlighting this peptide’s potential to modify the aggregation pathway of both Aβ40 and Aβ42. This distribution indicates that HP1 generates divergent combinations of kinetic acceleration and end-point epitope accessibility depending on the Aβ isoform used in the first seeding step. The homotypic Aβ42 control (Aβ42-42) occupied the highest PC2 position, consistent with its relatively high viability compared with peptide-conditioned aggregated samples.

The global organization of samples in PCA space supports the view that Aβ aggregation is not determined only by the components present in the final reaction, but also by the Interaction History through which aggregates are generated and propagated (Figure 8). Thus, differences introduced during the first aggregation cycle remain detectable after propagation, in a common monomeric substrate, resulting in distinct kinetic, conformational, and functional properties.

## 3. Conclusions

Our results support a view in which exogenous amyloid-like sequences act not only as direct modulators of Aβ aggregation, but also as factors that shape what becomes propagated in subsequent cycles. In this context, the relevant variable is not simply whether an exogenous peptide is present, but the Interaction History through which aggregates are generated. Even when a second reaction is performed with the same Aβ42 monomer substrate, aggregates arising from different first-generation templates can display distinct structural, kinetic, and toxic properties, supporting the transmission of properties across aggregation cycles. Whether this effect reflects purely conformational templating or also involves the presence of the exogenous peptide within the aggregates remains unresolved.

Within an energy landscape model of amyloid formation, multiple local minima represent distinct accessible conformational states, and early templating events can shift the probability of accessing one state over another (Figure 9). Exogenous sequences can therefore function as landscape modifiers that stabilize specific intermediates or favor specific growth pathways, producing strain-like outcomes whose properties can influence subsequent propagation. This interpretation is consistent with the structural heterogeneity of Aβ assemblies described in patient-derived material, where distinct fibril polymorphs and conformational variants have been associated with different amyloid subtypes and disease phenotypes [10,19,30,65,66].

**Figure 9.**
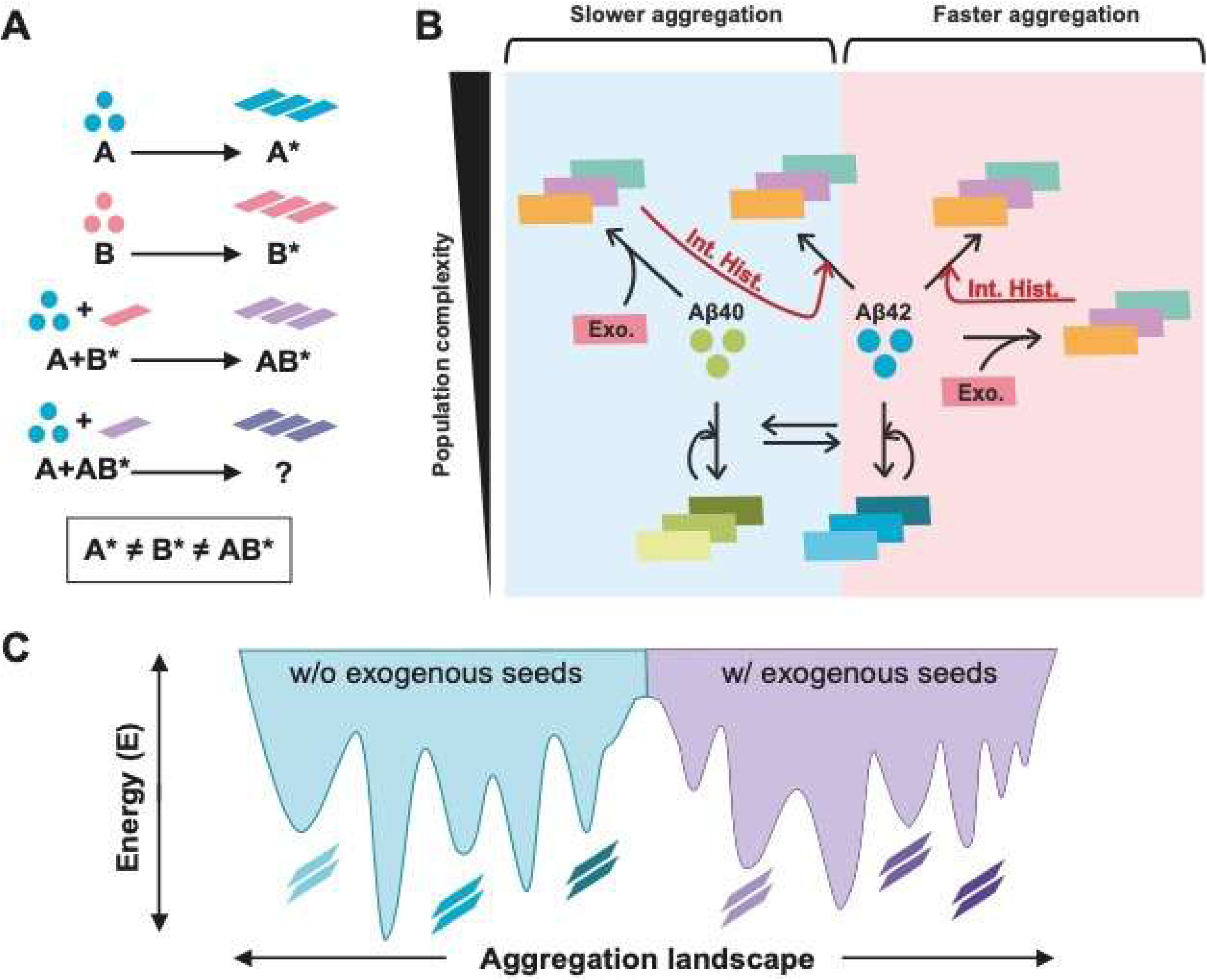
Conceptual model of Interaction History in Aβ aggregation and propagation. A) Schematic representation of how an exogenous amyloidogenic protein (B) may influence the aggregation of a host protein (A) across sequential aggregation events. A and B can independently form aggregates (A* and B*, respectively). When monomeric A aggregates in the presence of preformed B* aggregates, their interaction may generate an A-containing aggregate population (AB*) with properties that differ from those formed through homotypic aggregation. AB* can subsequently act as a seed for a new aggregation reaction involving monomeric A, whose outcome may therefore depend on the preceding molecular interactions. B) Conceptual representation of the multiple aggregation trajectories accessible to Aβ40 and Aβ42. Interactions with exogenous amyloid-like sequences (Exo.) and aggregates generated during previous aggregation events (Interaction History) may redirect these trajectories, producing differences in aggregation kinetics (slower to faster aggregation) and potentially increasing the diversity of accessible aggregate populations. C) Energy-landscape representation of this model. In the absence of exogenous interactions, protein assemblies populate a defined set of accessible states (blue). Interactions with exogenous sequences may reshape the landscape (purple), altering the relative accessibility of aggregate states and expanding the range of possible aggregation outcomes.

The PCA supports this concept by showing that Aβ40- and Aβ42-derived samples separate when structural, kinetic, and toxicity measurements are considered together (Figure 8). This suggests that the initial Aβ template constrains the range of outcomes accessible in the next propagation cycle. In this context, Aβ40-derived seeds appear to generate a narrower and less toxic set of aggregates, whereas Aβ42-derived histories allow a broader range of structural and toxic outcomes (Figure 8).

Overall, these results do not point to a single linear mechanism. Instead, they indicate that successive aggregation events are coupled: the molecular state generated in one cycle becomes the input for the next, and therefore influences the range of outcomes accessible during subsequent propagation. From this perspective, Aβ aggregation behaves as a history-dependent molecular process rather than as a series of independent reactions. Single-step aggregation assays may therefore underestimate aggregate diversity that emerges only when previous molecular interactions and successive propagation events are considered together.

Our findings are based on a controlled *in vitro* system that examines a defined set of molecular interactions and therefore does not reproduce the full complexity of Aβ aggregation *in vivo*. Further work will require higher-resolution structural and compositional analyses to define the molecular differences between the propagated assemblies and determine whether exogenous peptides remain associated with them or act only during earlier aggregation steps. Taken together, an Interaction History perspective provides a mechanistic rationale for how previous molecular interactions can shape the current amyloid state and thereby influence its subsequent behavior.

## 4. Methods

### 4.1. Protein alignment and net charge

Sequence alignments of the ten amyloid-forming core candidates were generated using Kalign implemented in Unipro UGENE v43.0 [67]. UGENE was also used for sequence visualization. Net charge at pH 7.0 was calculated using Protein Calculator v3.4 (https://protcalc.sourceforge.net/).

### 4.2. Peptide Preparation

All samples were prepared in low protein-binding microcentrifuge tubes (ThermoFisher Scientific, Waltham, MA, USA). The bacterial amyloid sequences were obtained in lyophilized form from the Peptide Synthesis Facility, Department of Experimental and Health Sciences, Universitat Pompeu Fabra (UPF). Peptides were initially solubilized in 1,1,1,3,3,3-hexafluoroisopropanol (HFIP), then divided into aliquots and dried overnight in a fume hood at room temperature. For the amyloid aggregation, we employed the same conditions as previously described [39]. In brief, the peptides were dissolved in DMSO to maintain monomeric states, followed by dilution in 50 mM phosphate buffer (PB), pH 7.4. All first-generation seeded reactions, including the homotypic Aβ42 control, contained the same final DMSO concentration.

Synthetic Aβ40 and Aβ42 were purchased from GenScript (Rijswijk, Netherlands). Stock solutions were prepared by dissolving 1 mg of peptide in a final concentration of 250 μM in 20 mM sodium phosphate buffer with 0.04% NH₃ and NaOH to achieve a pH of 11. The absence of aggregates was assessed using dynamic light scattering [39,68]. The peptide solution was then sonicated for 10 minutes (Fisherbrand Pittsburgh, PA, USA, FB15051) without sweep mode and stored at −80°C.

For the first-generation aggregation reaction, monomeric Aβ was diluted to a final concentration of 25 μM in 20 mM sodium phosphate buffer containing 100 mM NaCl, pH 7.4, in the presence or absence of pre-aggregated bacterial peptides. The mixtures were then incubated at 25°C for 24 h to generate the first-generation aggregates to be analyzed or used as seeds in the subsequent aggregation reactions. For the second-generation reaction, the resulting first-generation aggregates were used as seeds at a final concentration of 2.5 μM in fresh Aβ42 monomer (25 μM).

### 4.3. Thioflavin-T Binding and Aggregation Kinetics

Thioflavin-T (ThT) was dissolved in Milli-Q water to a concentration of 5 mM, filtered through a 0.2-μm filter, diluted to 0.5 mM, and stored at −20°C. ThT fluorescence during aggregation kinetics was measured every 10 min using a 440-nm excitation filter and a 480-nm emission filter on a plate reader (TECAN Infinite+ NANO) with bottom optics. Samples were prepared in flat-bottom, black, non-binding 96-well plates (Greiner Bio-One), with 100 μL of sample added per well. Each condition was measured in triplicate.

For seeding experiments, Aβ40 and Aβ42 peptide stocks at pH 11 were diluted to a final concentration of 25 μM in 20 mM sodium phosphate buffer with 100 mM NaCl, pH 7.4. The corresponding pre-aggregated peptides were sonicated for 5 minutes before being added to a final concentration of 2.5 μM. Control conditions included samples without Aβ42 but with the peptide, samples without Aβ42, and samples without peptides, all prepared with equivalent volumes of the corresponding buffers. All samples and controls were analyzed in three independent experiments, with three technical replicates per experiment. The pH in each well was measured at the beginning and end of the experiment to confirm stability throughout the aggregation process and to verify consistency across conditions. ThT was added to a final concentration of 20 μM. Aggregation reactions were conducted at 37°C without agitation.

The t₁/₂, defined as the time to reach 50% of the final fluorescence intensity, was measured for each condition. The lag phase was calculated as the time to reach 10% of the final fluorescence intensity. Also, the data were fitted to a sigmoidal curve using the Hill function:

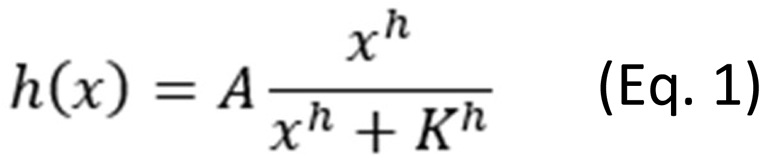

In short, A, h and K are the fitting parameters. The fittings were used to calculate the slope of the exponential phase of the aggregation kinetics, as done by Rupert et al. [20].

### 4.4. Transmission Electron Microscopy (TEM)

To prepare samples for TEM analysis, a 10 μL aliquot of the aggregated peptide solution (25 μM, incubated at 37°C for 24 hours without agitation) was applied to carbon-coated copper grids and allowed to adhere for 1 minute. Liquid excess was carefully blotted with Whatman filter paper [39]. Grids were then rinsed with distilled water and subsequently stained with 2% (w/v) uranyl acetate for 1 minute. After staining, grids were air-dried. Micrographs were obtained using a JEM-1400 transmission electron microscope (JEOL, Tokyo, Japan) set to an accelerating voltage of 80 keV.

### 4.5. Circular Dichroism Spectroscopy

Protein secondary-structure profiles of the final aggregated material were analyzed using far-UV circular dichroism (CD) spectroscopy in aggregation buffer (20 mM sodium phosphate buffer with 100 mM NaCl, pH 7.4). Spectra were recorded from 260 to 190 nm on a Jasco J-715 spectropolarimeter. Measurements were performed in 0.2 cm path-length quartz cuvettes, averaging 20 scans per sample at a scanning speed of 50 nm/min with a response time of 2 s. Spectra were baseline-corrected against the corresponding buffer and deconvoluted using the BeStSel web server to assess differences in secondary-structure composition across conditions.

### 4.6. Dot blot

For dot-blot analysis, 2 µL of each aggregate sample was spotted onto nitrocellulose membranes. This spotting step was repeated three times per sample to increase the total protein load. Membranes were allowed to air-dry for 5 min and subsequently blocked with 4% (w/v) bovine serum albumin (BSA).

Membranes were incubated with the following primary antibodies: the anti-oligomer antibody A11 (Invitrogen, AHB0052), the anti-amyloid fibrils polyclonal antibody (Invitrogen, PA5-77843), and the anti–amyloid-β N-terminal antibody 6E10 (epitope 3–8; chimeric rabbit monoclonal, Invitrogen MA5-48043). After washing, membranes were incubated with a goat anti-rabbit horseradish peroxidase (HRP)-conjugated secondary antibody (Bio-Rad; 1:3000). Immunoreactivity was visualized using enhanced chemiluminescence (ECL; Thermo Scientific) and images were acquired using a ChemiDoc MP Imaging System (Bio-Rad).

Dot-blot analyses were performed in three independent experiments and spot intensities were quantified using ImageJ. All conditions from the same experimental replicate were analyzed on the same membrane, and signal intensities were normalized to the unseeded Aβ42 condition on that membrane. For direct comparison of Aβ42- and Aβ40-derived seeding histories, normalized signals from matched conditions within each experimental replicate were expressed as Aβ42/Aβ40 ratios and subsequently ln-transformed.

### 4.7. Cell cytotoxicity assay

SH-SY5Y human neuroblastoma cells were cultured in Dulbecco’s modified Eagle’s medium/F-12 supplemented with GlutaMAX (DMEM/F-12 GlutaMAX), 10% (v/v) heat-inactivated fetal bovine serum (FBS), and 1% (v/v) penicillin/streptomycin. Cells were maintained at 37 °C in a humidified atmosphere containing 5% CO₂ and routinely grown in 75-cm² culture flasks.

Twenty-four hours after plating, neuronal differentiation was initiated by replacing the maintenance medium with differentiation medium and continued for 7 days, with medium renewal every 72 h. The differentiation medium consisted of DMEM/F-12 GlutaMAX supplemented with 2.5% (v/v) heat-inactivated FBS and 10 μM retinoic acid. Differentiation was monitored by microscopy based on morphological criteria.

For cytotoxicity assays, differentiated cells were seeded in 96-well plates at a density of 1 × 10⁴ cells per well. Preformed Aβ aggregates obtained at the end of the two-step aggregation protocol were added at a final concentration of 10 μM (monomer-equivalent). After 24 h of incubation, cell viability was assessed using the MTT assay. Briefly, the culture medium was removed and replaced with 100 μL of fresh medium containing 10 μL of MTT solution (5 mg/mL). Cells were incubated at 37 °C for 4 h, after which the supernatant was discarded, and 150 μL of dimethyl sulfoxide (DMSO) was added to dissolve the formazan crystals. Absorbance was measured at 580 nm using a TECAN Spark plate reader. Untreated control cells were incubated with the same buffer used for aggregate preparation, but without the addition of aggregates.

### 4.8. Principal component analysis

Principal component analysis (PCA) was performed using GraphPad Prism 9 to integrate kinetic, conformational, and toxicity measurements obtained from the two-step aggregation assays. Seven variables were included: lag phase, slope, half-time, anti-N-terminal, anti-oligomer and anti-fibril reactivity, and cell viability measured by MTT. The analysis comprised 12 end-point Aβ42 samples representing the different first-generation seeding conditions. Before PCA, each variable was standardized to a mean of 0 and a standard deviation of 1. The two principal components with the largest eigenvalues were selected for visualization. Sample scores and variable loadings for PC1 and PC2 were used to examine the relationships among aggregation histories and measured properties.

### 4.9. Statistics

Statistical analyses were performed using GraphPad Prism 9. Unless otherwise indicated, comparisons were performed using two-tailed unpaired t-tests. For experiments involving multiple peptide-wise comparisons, the false discovery rate (FDR) was controlled using the two-stage step-up method of Benjamini, Krieger and Yekutieli (Q = 0.05). The specific statistical tests used are indicated in the corresponding figure legends. Data are presented as mean ± SD unless otherwise stated.

## Declaration of Generative AI and AI-assisted technologies in the writing process

During the preparation of this work, the authors used ChatGPT (OpenAI) to assist with language editing and improve the clarity of the manuscript. The authors reviewed and edited the content and take full responsibility for the content of the manuscript.

## Acknowledgments

We thank Jacob Rupert for his comments and in-depth discussions, which provided valuable insights into this work. Transmission electron microscopy (TEM) images were acquired at the Servei de Microscòpia i Difracció de Raigs X (SMiDRX) of the Universitat Autònoma de Barcelona (UAB).

## Bibliography

[1] L.E. Hebert, P.A. Scherr, J.L. Bienias, D.A. Bennett, D.A. Evans, Alzheimer disease in the US population: prevalence estimates using the 2000 census, Arch. Neurol. 60 (2003) 1119–1122. 10.1001/ARCHNEUR.60.8.1119.

[2] F. Chiti, C.M. Dobson, Protein Misfolding, Amyloid Formation, and Human Disease: A Summary of Progress Over the Last Decade, Annu. Rev. Biochem. 86 (2017) 27–68. 10.1146/ANNUREV-BIOCHEM-061516-045115.

[3] F. Chiti, C.M. Dobson, Protein misfolding, functional amyloid, and human disease, Annu. Rev. Biochem. 75 (2006) 333–366. 10.1146/ANNUREV.BIOCHEM.75.101304.123901.

[4] T. Guo, D. Zhang, Y. Zeng, T.Y. Huang, H. Xu, Y. Zhao, Molecular and cellular mechanisms underlying the pathogenesis of Alzheimer’s disease, Mol. Neurodegener. 15 (2020). 10.1186/S13024-020-00391-7.

[5] H. Hampel, J. Hardy, K. Blennow, C. Chen, G. Perry, S.H. Kim, V.L. Villemagne, P. Aisen, M. Vendruscolo, T. Iwatsubo, C.L. Masters, M. Cho, L. Lannfelt, J.L. Cummings, A. Vergallo, The Amyloid-β Pathway in Alzheimer’s Disease, Mol. Psychiatry 26 (2021) 5481–5503. 10.1038/S41380-021-01249-0.

[6] G.F. Chen, T.H. Xu, Y. Yan, Y.R. Zhou, Y. Jiang, K. Melcher, H.E. Xu, Amyloid beta: structure, biology and structure-based therapeutic development, Acta Pharmacol. Sin. 38 (2017) 1205–1235. 10.1038/APS.2017.28.

[7] A.I. Sulatskaya, G.N. Rychkov, M.I. Sulatsky, E. V. Mikhailova, N.M. Melnikova, V.S. Andozhskaya, I.M. Kuznetsova, K.K. Turoverov, New Evidence on a Distinction between Aβ40 and Aβ42 Amyloids: Thioflavin T Binding Modes, Clustering Tendency, Degradation Resistance, and Cross-Seeding, Int. J. Mol. Sci. 23 (2022). 10.3390/IJMS23105513.

[8] J. Tran, D. Chang, F. Hsu, H. Wang, Z. Guo, Cross-seeding between Aâ40 and Aâ42 in Alzheimer’s disease, FEBS Lett. 591 (2017) 177–185. 10.1002/1873-3468.12526.

[9] L. Gu, Z. Guo, Alzheimer’s Aâ42 and Aâ40 peptides form interlaced amyloid fibrils, J. Neurochem. 126 (2013) 305–311. 10.1111/JNC.12202.

[10] Y. Yang, D. Arseni, W. Zhang, M. Huang, S. Lövestam, M. Schweighauser, A. Kotecha, A.G. Murzin, S.Y. Peak-Chew, J. MacDonald, I. Lavenir, H.J. Garringer, E. Gelpi, K.L. Newell, G.G. Kovacs, R. Vidal, B. Ghetti, B. Ryskeldi-Falco, S.H.W. Scheres, M. Goedert, Cryo-EM structures of amyloid-β 42 filaments from human brains, Science 375 (2022) 167–172. 10.1126/SCIENCE.ABM7285.

[11] G.A. Braun, A.J. Dear, K. Sanagavarapu, H. Zetterberg, S. Linse, Amyloid-â peptide 37, 38 and 40 individually and cooperatively inhibit amyloid-â 42 aggregation, Chem. Sci. 13 (2022) 2423–2439. 10.1039/D1SC02990H.

[12] C.N. Suire, S.O. Abdul-Hay, T. Sahara, D. Kang, M.K. Brizuela, P. Saftig, D.W. Dickson, T.L. Rosenberry, M.A. Leissring, Cathepsin D regulates cerebral Aβ42/40 ratios via differential degradation of Aβ42 and Aβ40, Alzheimers. Res. Ther. 12 (2020). 10.1186/S13195-020-00649-8.

[13] S.S. Kwak, K.J. Washicosky, E. Brand, D. von Maydell, J. Aronson, S. Kim, D.E. Capen, M. Cetinbas, R. Sadreyev, S. Ning, E. Bylykbashi, W. Xia, S.L. Wagner, S.H. Choi, R.E. Tanzi, D.Y. Kim, Amyloid-β42/40 ratio drives tau pathology in 3D human neural cell culture models of Alzheimer’s disease, Nat. Commun. 11 (2020). 10.1038/S41467-020-15120-3.

[14] M. Lee, W.M. Yau, J.M. Louis, R. Tycko, Structures of brain-derived 42-residue amyloid-â fibril polymorphs with unusual molecular conformations and intermolecular interactions, Proc. Natl. Acad. Sci. U. S. A. 120 (2023). 10.1073/PNAS.2218831120.

[15] F. Xiong, W. Ge, C. Ma, Quantitative proteomics reveals distinct composition of amyloid plaques in Alzheimer’s disease, Alzheimers. Dement. 15 (2019) 429–440. 10.1016/J.JALZ.2018.10.006.

[16] P. Bharadwaj, T. Solomon, B.R. Sahoo, K. Ignasiak, S. Gaskin, J. Rowles, G. Verdile, M.J. Howard, C.S. Bond, A. Ramamoorthy, R.N. Martins, P. Newsholme, Amylin and beta amyloid proteins interact to form amorphous heterocomplexes with enhanced toxicity in neuronal cells, Sci. Rep. 10 (2020). 10.1038/S41598-020-66602-9.

[17] K.H. Shim, M.J. Kang, Y.C. Youn, S.S.A. An, S.Y. Kim, Alpha-synuclein: a pathological factor with Aâ and tau and biomarker in Alzheimer’s disease, Alzheimers. Res. Ther. 14 (2022). 10.1186/S13195-022-01150-0.

[18] L.K. Clinton, M. Blurton-Jones, K. Myczek, J.Q. Trojanowski, F.M. LaFerla, Synergistic Interactions between Abeta, tau, and alpha-synuclein: acceleration of neuropathology and cognitive decline, J. Neurosci. 30 (2010) 7281–7289. 10.1523/JNEUROSCI.0490-10.2010.

[19] M.I. Ivanova, Y. Lin, Y.H. Lee, J. Zheng, A. Ramamoorthy, Biophysical processes underlying cross-seeding in amyloid aggregation and implications in amyloid pathology, Biophys. Chem. 269 (2021). 10.1016/J.BPC.2020.106507.

[20] J. Rupert, M. Monti, E. Zacco, G.G. Tartaglia, RNA sequestration driven by amyloid formation: the alpha synuclein case, Nucleic Acids Res. 51 (2023) 11466–11478. 10.1093/NAR/GKAD857.

[21] T. Janas, K. Sapoñ, M.H.B. Stowell, T. Janas, Selection of Membrane RNA Aptamers to Amyloid Beta Peptide: Implications for Exosome-Based Antioxidant Strategies, Int. J. Mol. Sci. 20 (2019). 10.3390/IJMS20020299.

[22] E. Álvarez-Marimon, H. Castillo-Michel, J. Reyes-Herrera, J. Seira, E. Aso, M. Carmona, I. Ferrer, J. Cladera, N. Benseny-Cases, Synchrotron X-ray Fluorescence and FTIR Signatures for Amyloid Fibrillary and Nonfibrillary Plaques, ACS Chem. Neurosci. 12 (2021) 1961–1971. 10.1021/acschemneuro.1c00048.

[23] A. Abelein, Metal Binding of Alzheimer’s Amyloid-â and Its Effect on Peptide Self-Assembly, Acc. Chem. Res. 56 (2023) 2653–2663. 10.1021/ACS.ACCOUNTS.3C00370.

[24] A.S. Pithadia, M.H. Lim, Metal-associated amyloid-â species in Alzheimer’s disease, Curr. Opin. Chem. Biol. 16 (2012) 67–73. 10.1016/J.CBPA.2012.01.016.

[25] J. Tittelmeier, S. Druffel-Augustin, A. Alik, R. Melki, C. Nussbaum-Krammer, Dissecting aggregation and seeding dynamics of á-Syn polymorphs using the phasor approach to FLIM, Commun. Biol. 5 (2022). 10.1038/S42003-022-04289-6.

[26] J. Seira Curto, M.R. Fernandez, J. Cladera, N. Benseny-Cases, N. Sanchez de Groot, Aâ40 Aggregation under Changeable Conditions, Int. J. Mol. Sci. 24 (2023). 10.3390/ijms24098408.

[27] R. Tycko, Amyloid polymorphism: structural basis and neurobiological relevance, Neuron 86 (2015) 632–645. 10.1016/J.NEURON.2015.03.017.

[28] L.D. Aubrey, S.E. Radford, How is the Amyloid Fold Built? Polymorphism and the Microscopic Mechanisms of Fibril Assembly, J. Mol. Biol. (2025) 169008. 10.1016/j.jmb.2025.169008.

[29] M. Wilkinson, Y. Xu, D. Thacker, A.I.P. Taylor, D.G. Fisher, R.U. Gallardo, S.E. Radford, N.A. Ranson, Structural evolution of fibril polymorphs during amyloid assembly, Cell 186 (2023) 5798–5811.e26. 10.1016/j.cell.2023.11.025.

[30] C. Duran-Aniotz, I. Moreno-Gonzalez, N. Gamez, N. Perez-Urrutia, L. Vegas-Gomez, C. Soto, R. Morales, Amyloid pathology arrangements in Alzheimer’s disease brains modulate in vivo seeding capability, Acta Neuropathol. Commun. 9 (2021). 10.1186/S40478-021-01155-0.

[31] J. Zhang, Y. Zhang, J. Wang, Y. Xia, J. Zhang, L. Chen, Recent advances in Alzheimer’s disease: Mechanisms, clinical trials and new drug development strategies, Signal Transduct. Target. Ther. 9 (2024). 10.1038/S41392-024-01911-3.

[32] J. Seira Curto, A. Surroca Lopez, M. Casals Sanchez, I. Tic, M.R. Fernandez Gallegos, N. Sanchez de Groot, Microbiome Impact on Amyloidogenesis, Front. Mol. Biosci. 9 (2022). 10.3389/FMOLB.2022.926702.

[33] Y. Belkaid, T.W. Hand, Role of the microbiota in immunity and inflammation, Cell 157 (2014) 121–141. 10.1016/J.CELL.2014.03.011.

[34] S. Holmqvist, O. Chutna, L. Bousset, P. Aldrin-Kirk, W. Li, T. Björklund, Z.Y. Wang, L. Roybon, R. Melki, J.Y. Li, Direct evidence of Parkinson pathology spread from the gastrointestinal tract to the brain in rats, Acta Neuropathol. 128 (2014) 805–820. 10.1007/S00401-014-1343-6.

[35] S.K. Kim, R.B. Guevarra, Y.T. Kim, J. Kwon, H. Kim, J.H. Cho, H.B. Kim, J.H. Lee, Role of Probiotics in Human Gut Microbiome-Associated Diseases, J. Microbiol. Biotechnol. 29 (2019) 1335–1340. 10.4014/JMB.1906.06064.

[36] D.N. Villageliu, D.R. Samuelson, The Role of Bacterial Membrane Vesicles in Human Health and Disease, Front. Microbiol. 13 (2022) 828704. 10.3389/fmicb.2022.828704.

[37] A. Taglialegna, L. Matilla-Cuenca, P. Dorado-Morales, S. Navarro, S. Ventura, J.A. Garnett, I. Lasa, J. Valle, The biofilm-associated surface protein Esp of Enterococcus faecalis forms amyloid-like fibers, Npj Biofilms Microbiomes 6 (2020) 15. 10.1038/s41522-020-0125-2.

[38] K. Oliphant, E. Allen-Vercoe, Macronutrient metabolism by the human gut microbiome: major fermentation by-products and their impact on host health, Microbiome 7 (2019). 10.1186/S40168-019-0704-8.

[39] J. Seira Curto, A. Dominguez Martinez, G. Perez Collell, E. Barniol Simon, M. Romero Ruiz, B. Franco Bordés, P. Sotillo Sotillo, S. Villegas Hernandez, M.R. Fernandez, N. Sanchez de Groot, Exogenous prion-like proteins and their potential to trigger cognitive dysfunction, Mol. Syst. Biol. 21 (2025) 1004–1029. 10.1038/s44320-025-00114-4.

[40] A. Fernández-Calvet, L. Matilla-Cuenca, M. Izco, S. Navarro, M. Serrano, S. Ventura, J. Blesa, M. Herráiz, G. Alkorta-Aranburu, S. Galera, I. Ruiz de los Mozos, M.L. Mansego, A. Toledo-Arana, L. Alvarez-Erviti, J. Valle, Gut microbiota produces biofilm-associated amyloids with potential for neurodegeneration, Nat. Commun. 15 (2024) 4150. 10.1038/s41467-024-48309-x.

[41] L. Meng, C. Liu, M. Liu, J. Chen, C. Liu, Z. Zhang, G. Chen, Z. Zhang, The yeast protein Ure2p triggers Tau pathology in a mouse model of tauopathy, Cell Rep. 42 (2023) 113342. 10.1016/j.celrep.2023.113342.

[42] L. Meng, C. Liu, Y. Li, G. Chen, M. Xiong, T. Yu, L. Pan, X. Zhang, L. Zhou, T. Guo, X. Yuan, C. Liu, Z. Zhang, Z. Zhang, The yeast prion protein Sup35 initiates á-synuclein pathology in mouse models of Parkinson’s disease, Sci. Adv. 9 (2023). 10.1126/SCIADV.ADJ1092.

[43] K.W. Tipping, P. van Oosten-Hawle, E.W. Hewitt, S.E. Radford, Amyloid Fibres: Inert End-Stage Aggregates or Key Players in Disease?, Trends Biochem. Sci. 40 (2015) 719–727. 10.1016/j.tibs.2015.10.002.

[44] R. Sabate, F. Rousseau, J. Schymkowitz, S. Ventura, What makes a protein sequence a prion?, PLoS Comput. Biol. 11 (2015). 10.1371/JOURNAL.PCBI.1004013.

[45] I. Pallarès, V. Iglesias, S. Ventura, The Rho Termination Factor of Clostridium botulinum Contains a Prion-Like Domain with a Highly Amyloidogenic Core, Front. Microbiol. 6 (2016). 10.3389/FMICB.2015.01516.

[46] A. Esteras-Chopo, L. Serrano, M. López De La Paz, The amyloid stretch hypothesis: recruiting proteins toward the dark side, Proc. Natl. Acad. Sci. U. S. A. 102 (2005) 16672–16677. 10.1073/PNAS.0505905102.

[47] Y.H. Liao, Y.J. Chang, Y. Yoshiike, Y.C. Chang, Y.R. Chen, Negatively charged gold nanoparticles inhibit Alzheimer’s amyloid-â fibrillization, induce fibril dissociation, and mitigate neurotoxicity, Small 8 (2012) 3631–3639. 10.1002/SMLL.201201068.

[48] H.M. Chan, L. Xiao, K.M. Yeung, S.L. Ho, D. Zhao, W.H. Chan, H.W. Li, Effect of surface-functionalized nanoparticles on the elongation phase of beta-amyloid (1-40) fibrillogenesis, Biomaterials 33 (2012) 4443–4450. 10.1016/J.BIOMATERIALS.2012.03.024.

[49] H. Liu, B. Xie, X. Dong, L. Zhang, Y. Wang, F. Liu, Y. Sun, Negatively charged hydrophobic nanoparticles inhibit amyloid â-protein fibrillation: The presence of an optimal charge density, React. Funct. Polym. 103 (2016) 108–116. 10.1016/J.REACTFUNCTPOLYM.2016.04.003.

[50] P. Arosio, T.P.J. Knowles, S. Linse, On the lag phase in amyloid fibril formation, Phys. Chem. Chem. Phys. 17 (2015) 7606. 10.1039/C4CP05563B.

[51] S.I.A. Cohen, S. Linse, L.M. Luheshi, E. Hellstrand, D.A. White, L. Rajah, D.E. Otzen, M. Vendruscolo, C.M. Dobson, T.P.J. Knowles, Proliferation of amyloid-â42 aggregates occurs through a secondary nucleation mechanism, Proc. Natl. Acad. Sci. U. S. A. 110 (2013) 9758–9763. 10.1073/pnas.1218402110.

[52] M.R.H. Krebs, L.A. Morozova-Roche, K. Daniel, C. V. Robinson, C.M. Dobson, Observation of sequence specificity in the seeding of protein amyloid fibrils, Protein Sci. 13 (2004) 1933–1938. 10.1110/PS.04707004.

[53] K. Konstantoulea, P. Guerreiro, M. Ramakers, N. Louros, L.D. Aubrey, B. Houben, E. Michiels, M. De Vleeschouwer, Y. Lampi, L.F. Ribeiro, J. de Wit, W. Xue, J. Schymkowitz, F. Rousseau, Heterotypic Amyloid â interactions facilitate amyloid assembly and modify amyloid structure, EMBO J. 41 (2022). 10.15252/embj.2021108591.

[54] S. Subedi, S. Sasidharan, N. Nag, P. Saudagar, T. Tripathi, Amyloid Cross-Seeding: Mechanism, Implication, and Inhibition, Molecules 27 (2022). 10.3390/MOLECULES27061776.

[55] G. Meisl, X. Yang, E. Hellstrand, B. Frohm, J.B. Kirkegaard, S.I.A. Cohen, C.M. Dobson, S. Linse, T.P.J. Knowles, Differences in nucleation behavior underlie the contrasting aggregation kinetics of the Aβ40 and Aβ42 peptides, Proc. Natl. Acad. Sci. U. S. A. 111 (2014) 9384–9389. 10.1073/pnas.1401564111.

[56] N.M. Vogt, R.L. Kerby, K.A. Dill-McFarland, S.J. Harding, A.P. Merluzzi, S.C. Johnson, C.M. Carlsson, S. Asthana, H. Zetterberg, K. Blennow, B.B. Bendlin, F.E. Rey, Gut microbiome alterations in Alzheimer’s disease, Sci. Rep. 7 (2017). 10.1038/S41598-017-13601-Y.

[57] F. Panza, M. Lozupone, V. Solfrizzi, M. Watling, B.P. Imbimbo, Time to test antibacterial therapy in Alzheimer’s disease, Brain 142 (2019) 2905–2929. 10.1093/BRAIN/AWZ244.

[58] C. Na, M. Kim, G. Kim, Y. Lin, Y.H. Lee, W. Bal, E. Nam, M.H. Lim, Distinct Aggregation Behavior of N-Terminally Truncated Aâ4-42 Over Aâ1-42 in the Presence of Zn(II), ACS Chem. Neurosci. 16 (2025) 732–744. 10.1021/ACSCHEMNEURO.4C00831.

[59] Y. Bouter, K. Dietrich, J.L. Wittnam, N. Rezaei-Ghaleh, T. Pillot, S. Papot-Couturier, T. Lefebvre, F. Sprenger, O. Wirths, M. Zweckstetter, T.A. Bayer, N-truncated amyloid â (Aâ) 4-42 forms stable aggregates and induces acute and long-lasting behavioral deficits, Acta Neuropathol. 126 (2013) 189–205. 10.1007/S00401-013-1129-2.

[60] M.A. Wälti, F. Ravotti, H. Arai, C.G. Glabe, J.S. Wall, A. Böckmann, P. Güntert, B.H. Meier, R. Riek, Atomic-resolution structure of a disease-relevant Aβ(1-42) amyloid fibril, Proc. Natl. Acad. Sci. U. S. A. 113 (2016) E4976–E4984. 10.1073/PNAS.1600749113.

[61] P. Arosio, R. Cukalevski, B. Frohm, T.P.J. Knowles, S. Linse, Quantification of the Concentration of Aâ42 Propagons during the Lag Phase by an Amyloid Chain Reaction Assay, J. Am. Chem. Soc. 136 (2014) 219–225. 10.1021/JA408765U.

[62] M. Wolff, D. Unuchek, B. Zhang, V. Gordeliy, D. Willbold, L. Nagel-Steger, Amyloid â Oligomeric Species Present in the Lag Phase of Amyloid Formation, PLoS One 10 (2015) e0127865. 10.1371/JOURNAL.PONE.0127865.

[63] A. Jan, O. Adolfsson, I. Allaman, A.L. Buccarello, P.J. Magistretti, A. Pfeifer, A. Muhs, H.A. Lashuel, Abeta42 neurotoxicity is mediated by ongoing nucleated polymerization process rather than by discrete Abeta42 species, J. Biol. Chem. 286 (2011) 8585–8596. 10.1074/JBC.M110.172411.

[64] C. Haass, D.J. Selkoe, Soluble protein oligomers in neurodegeneration: lessons from the Alzheimer’s amyloid beta-peptide, Nat. Rev. Mol. Cell Biol. 8 (2007) 101–112. 10.1038/NRM2101.

[65] J. Rasmussen, J. Mahler, N. Beschorner, S.A. Kaeser, L.M. Häsler, F. Baumann, S. Nyström, E. Portelius, K. Blennow, T. Lashley, N.C. Fox, D. Sepulveda-Falla, M. Glatzel, A.L. Oblak, B. Ghetti, K.P.R. Nilsson, P. Hammarström, M. Staufenbiel, L.C. Walker, M. Jucker, Amyloid polymorphisms constitute distinct clouds of conformational variants in different etiological subtypes of Alzheimer’s disease, Proc. Natl. Acad. Sci. U. S. A. 114 (2017) 13018–13023. 10.1073/PNAS.1713215114.

[66] P. Chaudhuri, K.P. Prajapati, B.G. Anand, K. Dubey, K. Kar, Amyloid cross-seeding raises new dimensions to understanding of amyloidogenesis mechanism, Ageing Res. Rev. 56 (2019). 10.1016/J.ARR.2019.100937.

[67] K. Okonechnikov, O. Golosova, M. Fursov, UGENE team, Unipro UGENE: a unified bioinformatics toolkit., Bioinformatics 28 (2012) 1166–7. 10.1093/bioinformatics/bts091.

[68] N. Benseny-Cases, O. Klementieva, J. Cladera, In vitro oligomerization and fibrillogenesis of Amyloid-beta peptides, Subcell. Biochem. 65 (2012) 53–74. 10.1007/978-94-007-5416-4_3.

